# Longitudinal modelling reveals widespread non-additive genetic effects underlying developmental plasticity

**DOI:** 10.64898/2025.12.19.695443

**Authors:** Ralph Porneso, Alexandra Havdahl, Espen Eilertsen, Eivind Ystrom

## Abstract

Developmental plasticity, or the capacity of genotypes to produce different phenotypes under varying contexts, is poorly characterized in humans. Here we developed a longitudinal variance and a genotype-by-time model able to disentangle genetic effects from between-subject variability. Applied to early growth (infant length and BMI) and cognitive traits (math and reading) of 45,000 to 65,000 individuals, our models discovered 76 lead putative plasticity loci. Nearly all loci are novel; six replicate signals from independent interaction studies. We found evidence that additive genetic variation captures marginal effects from interactions, challenging the assumption that effect estimates from genome-wide association studies (GWAS) are purely additive. Notably, over 90% of putative plasticity loci are involved in distal intra-chromatin interactions implicating regulatory activity. Our findings provide molecular evidence that non-additive genetic variation contributes to complex traits and highlight the utility of longitudinal models in uncovering the biological underpinnings of trait plasticity.

---

Developmental plasticity is the capacity of a genotype to give rise to different phenotypes across varying contexts, especially in early life or during critical periods^1^. It is an important property in biological systems because it allows individuals to adapt to their environment to increase fitness and survival. Despite its importance, the genetic basis of developmental plasticity remains poorly understood, in part because of technical challenges and limitations in traditional statistical methods, e.g. small effect sizes and increased multiple testing burden in pairwise interaction models.

Genome-wide association studies (GWAS) have been instrumental in uncovering biological mechanisms in disease and health. The additive effects estimated in this framework are marginal because they capture the effects of unmodeled variables. For example, GWAS estimates include the effects of other genetic loci in proximity to the measured locus. This implies that they also capture effects from undetected genetic interactions. This misspecification results in genotype-dependent residual variability, or genetic heteroscedasticity, which may be leveraged to test for interactions without explicit knowledge and measurement of the interacting variable. Variance-based approaches exploit this property as a heuristic to identify variance quantitative trait loci (vQTL). Loci involved in gene-environment or epistatic interactions generate unequal variances, resulting in dispersion of trait values in a genotype-dependent manner^2–8^. We refer to this as non-additive genetic effect throughout.

Variance-based approaches, however, have lower power than methods that explicitly model interactions and may miss loci with small non-additive effects^2,9,10^. Furthermore, estimates from population-based vQTL studies capture signal from between-subject variability, which may include non-genetic sources, such as selection and phantom vQTL^5,11^. A recent study reported that non-additive effects from within-individual and population-level studies are only moderately correlated (r=0.49)^12^, suggesting effect estimates from the latter may include environmental sources.

Longitudinal models can disentangle between- from within-individual variation and estimate genetic effects that are less confounded by environmental factors. Importantly, such approaches also allow genetic effects to vary over time. Age may represent environmental and biological exposures present at specific time points. By using age to proxy for an individual’s varying context during development, longitudinal models may be used to detect genetic interactions with factors that have age- or time-specific expression.

In this pre-registered study, we aim to identify loci associated with trait level and/or variability while minimizing influences from environmental factors (**Figure 1**). We fitted a log-linear variance model using a mixed effect framework that jointly estimates additive and non-additive genetic effects to longitudinal growth traits in ∼45,000 infants and cognitive traits in ∼65,000 early adolescents from the Norwegian Mother, Father and Child cohort study (MoBa). We assessed the performance of our model in silico and compared it to trajGWAS^11^, a pipeline that similarly estimates additive and non-additive genetic effects. Loci exhibiting time-varying effects may be involved in gene-environment and epistatic interaction. We fitted a genotype-by-time model to our traits as a complementary approach. This strategy circumvents the increased multiple testing burden in pairwise interaction models while addressing the limited power in variance-based approaches. Here, we demonstrate that a simpler longitudinal model is a viable approach to identify loci with non-additive effects, which may be involved in developmental plasticity.

**Figure 1.**
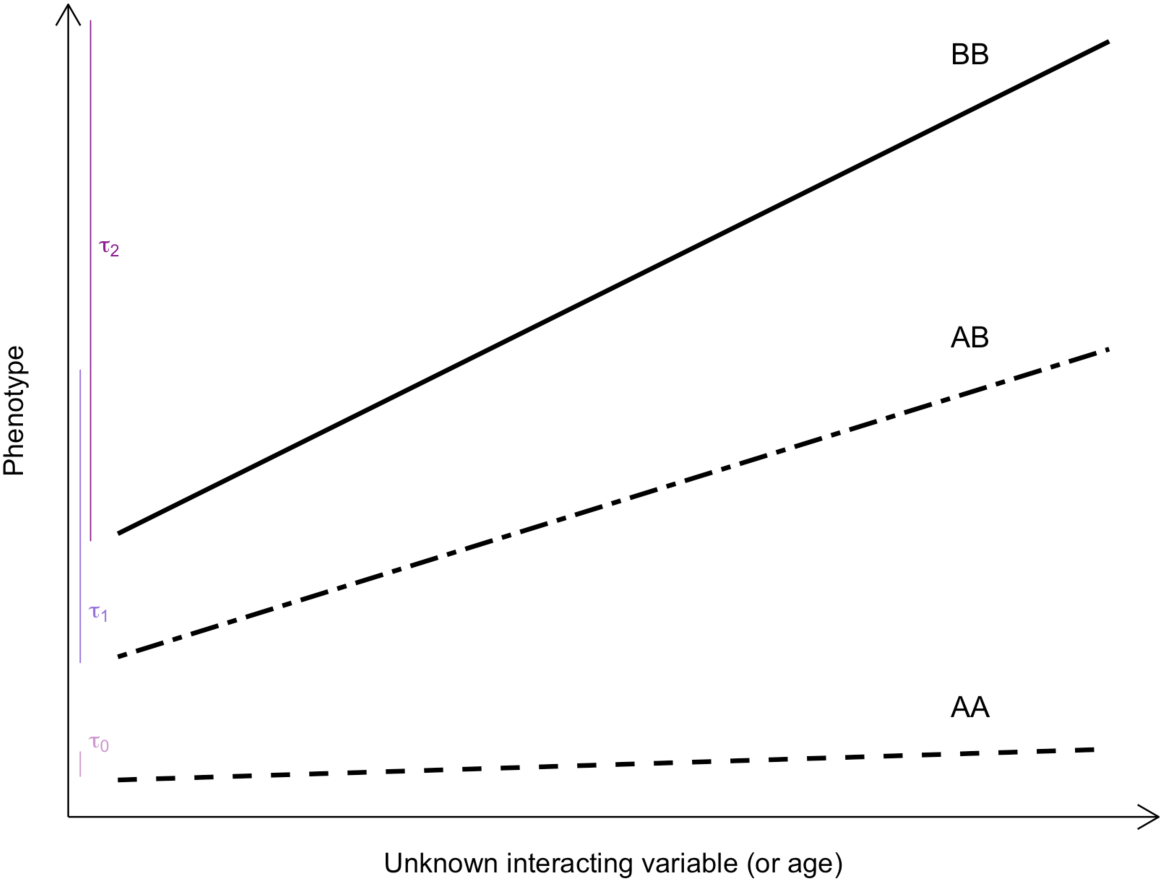
Longitudinal models can discover genetic interactions without pairwise interaction testing. A genotype interacting with an unknown factor increases the phenotype across factor and genotype levels, resulting in uneven phenotypic variability between groups. A longitudinal log-linear variance model infers interactions by allowing and estimating unequal variances across genotype groups net of between-subject variability, i.e. 𝛕_0_, 𝛕_1_, and 𝛕_2_ (purple). Non-additive effect is used throughout this paper when 𝛕_1_, 𝛕_2_ ≶ 𝛕_0_. In connection, a genotype-by-time interaction, or time-varying effect, may result when an unknown interacting variable is correlated with time or age. This is illustrated above as nonparallel lines across genotype groups. Non-additive and time-varying effects may represent undetected gene-gene or gene-environment interactions. As such, we refer to genotypes exhibiting these effects as putative plasticity loci.

## RESULTS

### Model performance

We evaluated bias, power, and false discovery rate in our variance model relative to trajGWAS under 3 scenarios (**Methods**). The first scenario involves a random intercept in the data generation process where trait levels differ between individuals. This is the simplest case where measures are collected from the same individual over multiple timepoints. Our expectation is for both models to perform well in this scenario. The second involves a data generating model with a random intercept and a random slope representing individual differences in developmental trajectories. Both our log-linear variance model and trajGWAS give the option to fit a random slope. However, fitting a random slope to our data with only 3 time points led to non-convergence. Scenario 2 aims to test if our model will produce biased estimates and increase false positives for failing to account for individual differences in developmental trajectories. The third and most complex scenario involves random intercepts in the mean and the residual. Here, trait levels and variability differ across individuals. TrajGWAS, designed for longitudinal analysis of phenotypes with 5 or more measures, fits a random intercept in the residual by default and is expected to perform well in this scenario.

Our log linear variance model provided unbiased estimates with narrower standard errors than trajGWAS across all scenarios (**Supplemental Figure 7**). Power was comparable between models. In terms of false discovery rate, our model retained the correct alpha level in scenarios 1 and 2 whereas trajGWAS’ was slightly inflated. This was reversed as expected in scenario 3, where our model showed minimal inflation (i.e. ∼7%) (**Supplemental Figure 6**). Given this, and because the true data generating process is difficult to discern in our dataset with only 3 time points, we fitted both models to our longitudinal traits.

### Putative plasticity loci exhibit intra-chromatin interactions and map to genes enriched during prenatal development in trait-relevant tissues

We fitted a log-linear variance model to our longitudinal traits using a mixed effect framework that accounts for between-subject variability, genetic relatedness, and population structure (**Methods)**. Across traits, we identified genetic loci associated with trait level and/or variability: 60 for infant length, 194 for BMI, 48 for math and 24 for reading (**Figure 2**). Standardized math and reading scores are heavily skewed (**Supplemental Figure 3**), which may drive the non-additive effects we observed for these traits. Inverse normal transformation attenuated effects of non-additive loci by 0.005 on average, with very minimal impact on the test statistics for math but not reading (**Supplemental Figure 12**).

**Figure 2.**
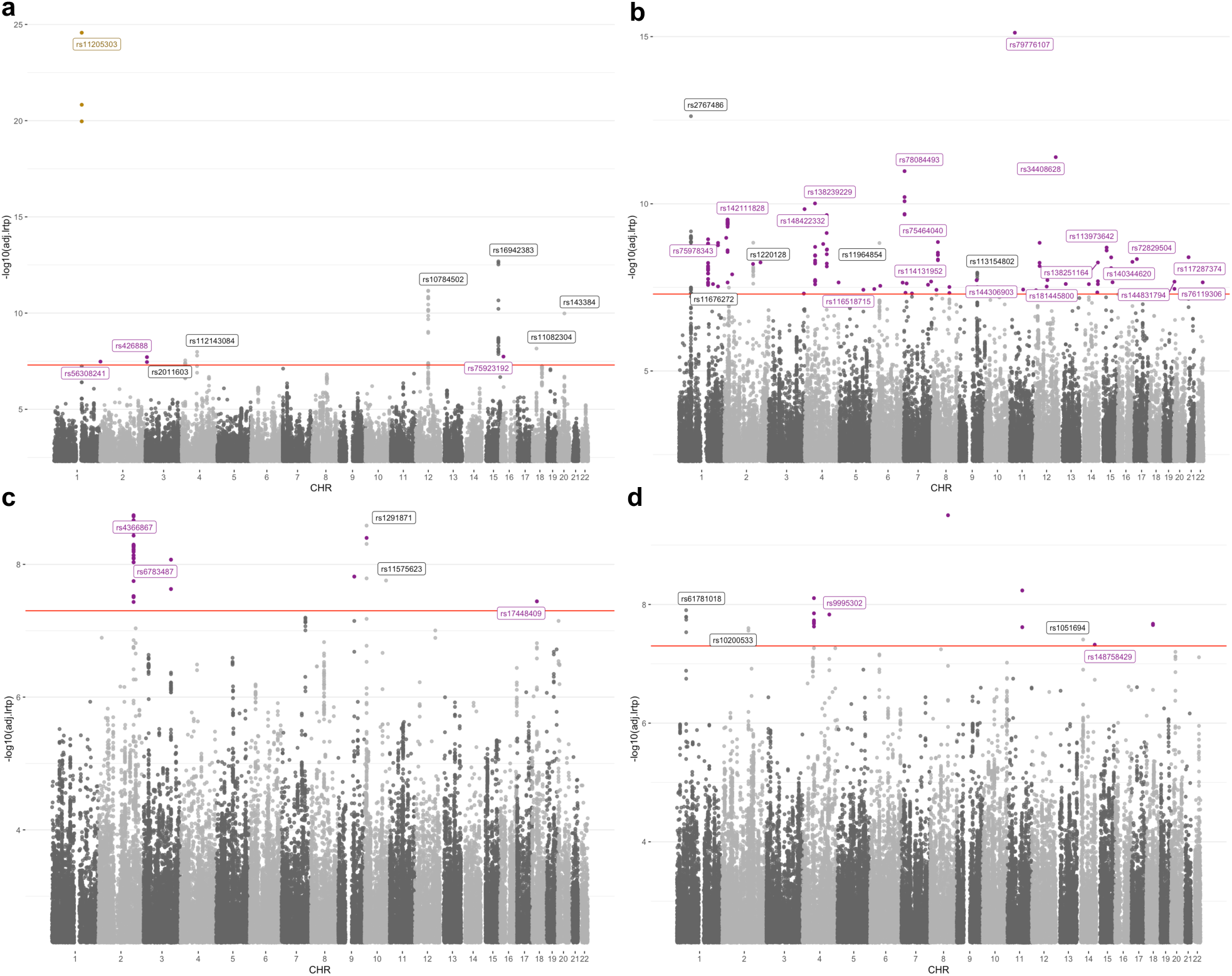
Putative plasticity loci. Manhattan plots for infant length (a), BMI (b), math (c), and reading (d). Lead loci are labeled and color-coded according to their effect on the trait: additive (grey), non-additive (purple), time-varying (gold). All test statistics were adjusted using genomic control. QQ plots are in Supplemental Figure 8. In (a), rs11205303 shows evidence of a time-varying effect (Table 1) and reported in ^17^ to be in suggestive epistatic interaction with rs17583662. In (b), only lead loci with the lowest p-value for each chromosome are labeled. Note that majority have MAF < 0.10 but with large effects detectable by our study (Supplemental Figures 9-10). In (c, d), only non-additive lead loci that survived inverse normal transformation are labeled.

**Table 1.**
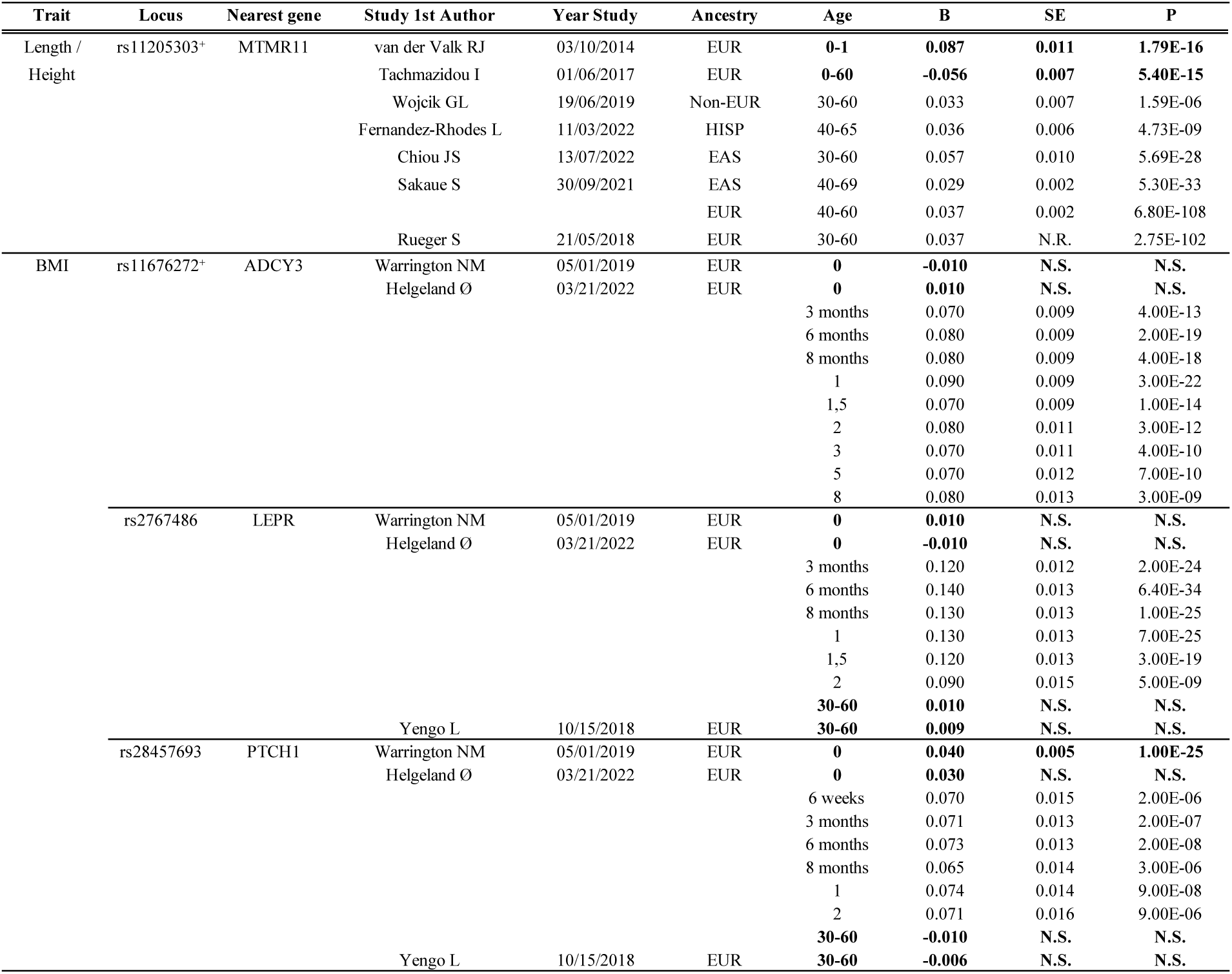
Additive loci with evidence of time-varying effects based on previous GWAS. Loci superscripted by a cross overlap with previous epistasis or trait change study. For infant length, rs11205303 is reported in ^17^ to be in suggestive epistatic interaction with a lncRNA variant. For BMI, rs11676272 has previous association with rate of BMI change in late childhood^17^. N.S. = not significant.

Except for infant length, 75% of the loci exhibit non-additive effects. Most are novel, suggesting they do not influence trait levels in general (**Supplemental Table 3**). In contrast, almost all additive loci have been tagged in prior GWAS (**Supplemental Table 8-13**).

To explore potential regulatory mechanisms, we functionally annotated all lead loci using FUMA^13^. A larger proportion of non-additive lead loci (62 of 70) than additive loci (9 of 17) are genome-wide significant for distal intra-chromatin interactions, implicating long-range regulatory activity as a potential contributor to trait plasticity (**Figure 3a-b**).

**Figure 3.**
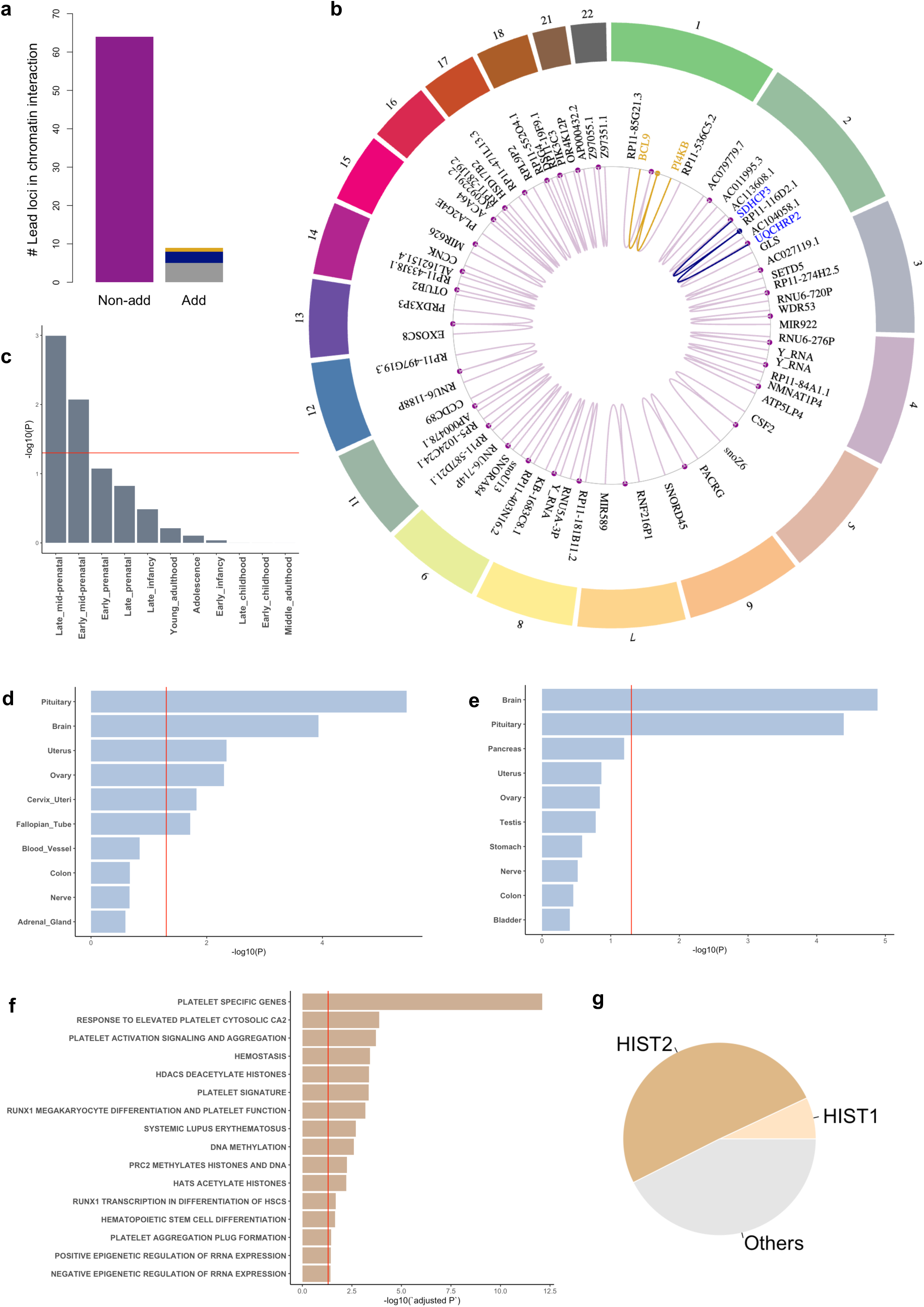
Putative plasticity loci exhibit distal intra-chromatin interactions implicating cis-regulatory mechanism and map to genes enriched during prenatal development in trait-relevant tissues. Non-additive and additive lead loci in intra-chromatin interactions; about half of the additive loci have or show evidence of time-varying effects (gold and blue, respectively) (a). Top lead loci (dots) and the enhancer or promoter region of the genes they interact with; rs11205303 (gold) and rs11676272 (blue) (b). Genes linked to math loci are enriched during prenatal development (c) and in brain tissues (d). Same as d but for reading (e). Infant length loci map to genes enriched in canonical pathways related to megakaryocytes and epigenetic regulation (f). Over half of the genes in these pathways belong to the HIST1/2 histone gene clusters (g). In (b), only the furthest gene up and downstream of the locus are shown. More comprehensive circos plots for each chromosome per trait are available in Supplemental Figures 19-20. Note that Y_RNA is a pseudogene found in multiple chromosomes.

We performed biological annotation to check if our hits tag genes enriched in trait-relevant tissues. Genes linked to infant length loci are differentially expressed in ovarian and uterine tissues as well as fetal and embryonic megakaryocytes, i.e. bone marrow cells involved in platelet production. They are enriched in pathways for hematopoietic stem cell differentiation and DNA methylation, with over half belonging to the canonical histone gene clusters, HIST1/2 (**Figure 3f-g**). Genes tagged by math and reading loci are also differentially expressed but in brain tissues, with math-related genes further enriched during mid-to-late prenatal brain development (**Figure 3c-e**).

These findings implicate prenatal developmental processes, and the lack of GWAS overlap with non-additive loci^14^, along with their enrichment for intra-chromatin interactions, suggests a role for cis-regulatory elements in early growth and cognitive traits.

### Concordant analytic results using trajGWAS

We re-ran our analyses in trajGWAS as a robustness check. TrajGWAS does not return effect estimates at the association testing step. Hence, we extracted the resulting Wald test p-value for comparison. This is equivalent to the joint effects p-value in our likelihood ratio test, which checks for the presence of additive and/or non-additive genetic effects.

The p-value correlations between our log linear variance model and trajGWAS are strong and significant (across traits p < 2 x 10^-16^): 0.83 for infant length, 0.75 for BMI, 0.67 for math, and 0.66 for reading (**Supplemental Figure 16**). They are not unity, however. One reason for this is that trajGWAS imputes missing genotypes. We took a more conservative approach by removing individuals with missing genotypes at each locus. Differences in the estimation procedure and model parameters are another reason. TrajGWAS uses a combination of Method of Moments (MoM) and M-asymptotic estimation for computational efficiency, whereas our model uses maximum likelihood. TrajGWAS provides less precise estimates compared to our approach partly due to these estimators^15^. In addition, trajGWAS fits additional random terms to the data, which further increases uncertainty in the estimates and contributes to more conservative p-values.

Given the above, we checked for overlapping hits between our approach and trajGWAS using the nominal genome-wide significance threshold of 5 x 10^-6^. We found 80% of the hits are shared between our log linear variance model and trajGWAS. Relaxing this threshold further to 5 x 10^-4^ increased the overlap to 96% (**Supplemental Tables 4-7**). This suggests both approaches capture the same genetic signal, and trajGWAS’ more conservative p-values are likely due to its estimators and the additional random terms in the residual it fits to the data as a default.

### Additive effects may reflect non-additive genetic variation from unmodelled interactions

Additive genetic effects are significantly correlated with non-additive and time-varying genetic effects within traits (**Figure 4**). This finding is consistent with theoretical derivations^16^ and our simulations, which demonstrates that epistasis induces both additive and time-varying genetic effects (**Supplemental Figure 1; Supplementary Note 1**).

**Figure 4.**
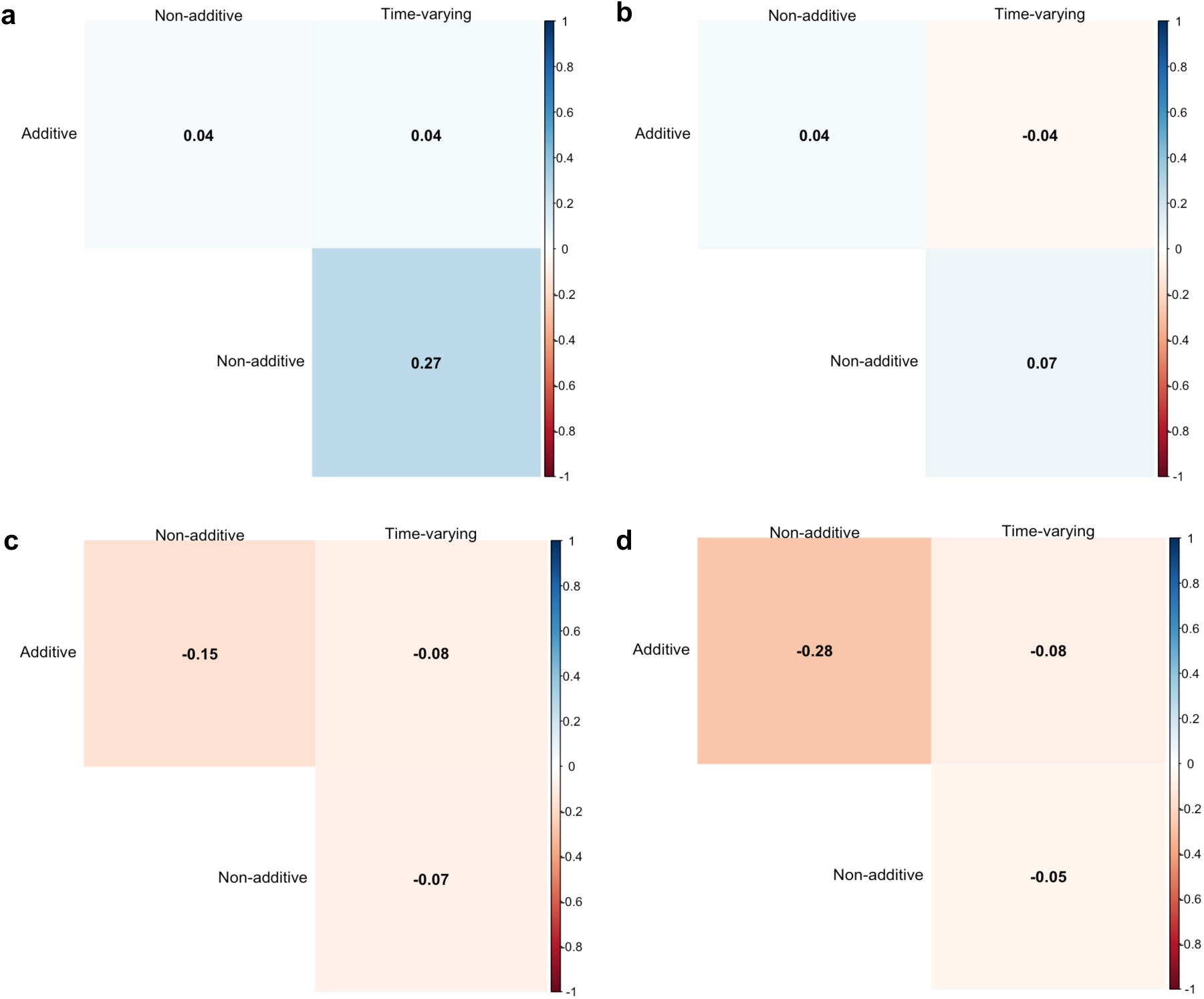
Additive, non-additive and time-varying effects are correlated. Correlation matrix for infant length (a), BMI (b), math (c), and reading (d). Effect sizes are centered and standardized by the minor allele frequency (MAF). All p-values < 2 x 10^-16^.

These within-trait correlations may have multiple interpretations. One is “effect pleiotropy”, where a locus affecting trait level also influences variability. A significant correlation may also arise from the general mean-variance relationship in non-normal traits like BMI^4,6^. Another possibility is that these correlations reflect marginal associations produced when interaction effects are absorbed into additive genetic estimates. To test whether these correlations reflect biologically meaningful interactions, we examined additive loci in our study for evidence of epistasis, non-additive and time-varying additive effects.

The top hit for infant length, rs11205303, a nonsynonymous mutation in exon 6 of MTMR11, has been reported in multiple height and birth length GWAS. **Table 1** shows its additive effect is consistent across ancestries but varies over time, suggesting that its marginal effect may reflect interactions with other factors. Supporting evidence comes from the largest epistasis study to date, which reported rs11205303 is in suggestive interaction with rs17583662, a long intergenic noncoding RNA (lncRNA) variant on chromosome 2, decreasing adult height by 0.37 mm^17^. Notably, the reported additive effect of rs11205303 is attenuated when an epistatic interaction is included in the model (**Supplemental Table 14**). We ran the same interaction model in our dataset but did not find an effect, suggesting that the time-varying effect of rs11205303 may be due to interaction with other factors during infancy.

Three BMI loci show a similar pattern, with rs11676272, a nonsynonymous variant in exon 2 of ADCY3 on chromosome 2, having a reported association with the rate of BMI change in late childhood^18^ (**Table 1-2**). GWAS of trait change have been used in the past to detect genetic association with trait variability^12^. These findings suggest that these loci are strong candidates for future pairwise interaction testing.

To rule out the potential role of linkage disequilibrium (LD) in driving the correlation between additive and non-additive effects, we retrieved all non-additive lead loci and extracted variants that are in LD with them. We compared this list against marginally significant additive loci. We did not find any additive loci in LD with non-additive lead loci in all traits.

Overall, these results indicate that additive effects estimated from GWAS may partly reflect non-additive genetic variation, implying that vQTL approaches prioritizing variants based solely on non-additive effects are likely to miss loci involved in interactions.

### Time-varying additive effects recapitulate genetic interactions in early growth traits using birth measures as baseline

Time-varying additive effects may reflect epistatic and gene-environment interactions, with time serving as a proxy for the changing context across the life course. Context can either be external or internal. External refers to environmental factors that an individual is exposed to, whereas internal encompasses biological and physiological processes and may include molecular switches and genes that are dynamically activated or silenced during development. We fitted a genotype-by-time model to our traits under the traditional GWAS framework while controlling for between-subject variability, genetic relatedness, and population structure (**Methods**).

Our model discovered loci with time-varying effects in all traits except for math (**Supplemental Figure 17**). We compared the effect estimates in this model with non-additive effect estimates from our variance model and assessed whether any of the loci overlapped between approaches. We found significant effect correlations across all traits (**Figure 4**). However, we did not find any locus that exhibits both time-varying and non-additive effects.

**Table 1** shows additive loci for infant length and BMI that exhibit differing effects at birth and/or in adulthood. To check if the lack of overlap may be due to the selected timepoints in our study, we re-ran our analyses using length and BMI measures at birth, 8, and 12 months (not pre-registered).

We corroborated that rs11205303 exhibits a highly significant time-varying effect on infant length along with two other loci in high LD with it: rs67807996 (r^2^ = 0.88) and rs11205277 (r^2^ = 0.84) (**Figure 5a**).

**Figure 5.**
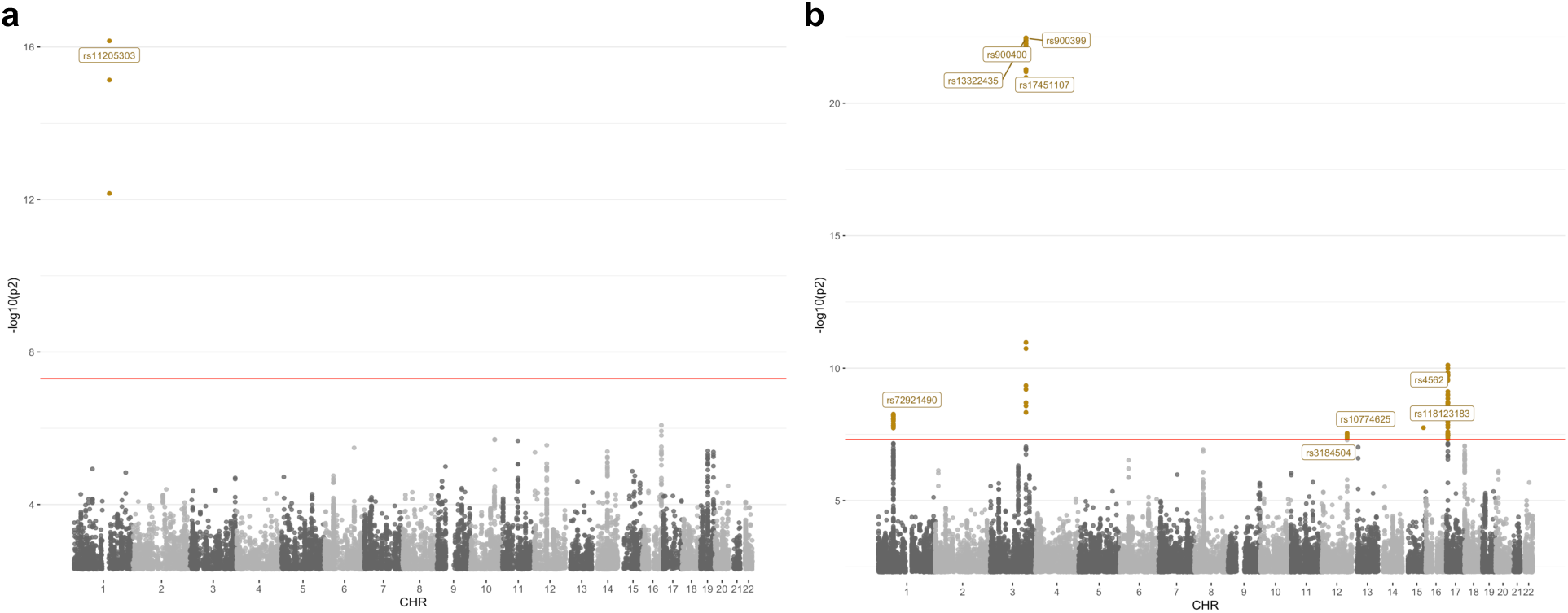
Loci with time-varying effects in early growth traits using birth measures as baseline. Genotype-by-time Manhattan plots for infant length (a) and BMI (b). Lead loci are labeled and color-coded as in Figure 2. We confirmed that rs11205303 exhibits a time-varying additive effect (a) and found four additional loci for BMI that overlap with previous epistasis and vQTL studies: rs900400, rs900399, rs17451107, and rs3184504 (b). QQ plots are in Supplemental Figure 13. All lead loci exhibit intra-chromatin interactions (Supplemental Figure 21).

For BMI, we found four additional loci that overlap with previous epistasis and vQTL studies. rs3184504, a nonsynonymous variant in exon 2 of SH2B3 on the q-arm of chromosome 12, was reported in epistatic interaction with rs7970953, an intronic variant in SOX5 on the p-arm of the same chromosome^19^. While rs900400, rs900399, and rs17451107 are lncRNA variants previously reported in population-level BMI and fat percentage vQTL studies, where they were shown in putative interaction with diet and/or level of physical activity^6,8,12^ (**Table 2**; **Figure 5b**).

**Table 2.**
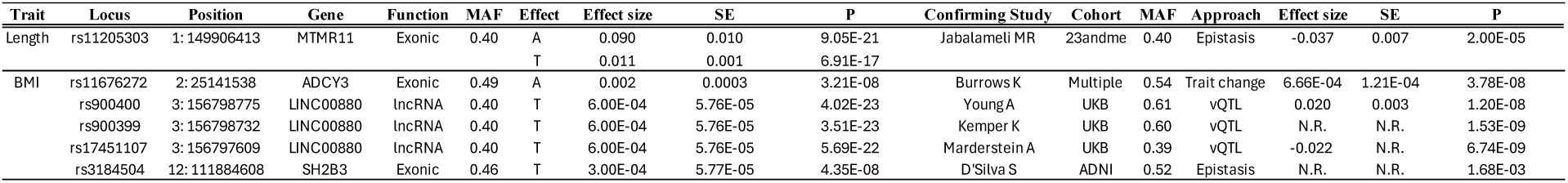
Putative plasticity loci overlapping with epistasis and vQTL studies. Coding and noncoding variants with additive and/or time-varying effects are implicated in genetic interactions. Effect sizes are standardized by MAF. Exonic variants in MTMR11 and SH2B3 are reported to be in epistatic interaction with noncoding variants in RPL34-DT and the intronic region of SOX5, respectively. Introns may function as regulatory elements^46^. Multiple cohorts in Burrows K include ALSPAC, CHOP, NFBC1966, NFBC1986, and OBE. A = Additive, T = Time-varying; N.R. = not reported.

These results illustrate that modelling time-varying effects using developmentally relevant time points can recover known interacting loci from epistasis and cross-sectional vQTL studies.

## DISCUSSION

Longitudinal models offer the temporal resolution needed to detect and disentangle the distinct and dynamic effects of common genetic variants on complex traits. We showed in simulations that genetic interactions may underlie these effects, which our empirical results supported, with several loci that were independently reported as vQTLs or interacting with distal loci in large-scale studies. This convergence across multiple cohorts and modeling frameworks indicates that our approach captures genuine genetic signals of developmental plasticity.

Functional genomics and epidemiological studies on trait-related diseases contextualize some of the highlighted loci in our study (**Table 2**). For example, ADCY3 is a key component of the hypothalamic signaling pathway involved in energy homeostasis and shows age-dependent influences in early childhood^20^. Consistently, mutations in ADCY3 have been shown to cause a severe form of monogenic obesity, where a homozygous exonic mutation in mice results in the protein product’s loss of catalytic activity^21^. Similarly, SH2B3 has been shown to increase the likelihood of coronary heart disease. Individuals homozygous for rs3184504 who take statins show a 48% relative risk reduction for the disease^22^. These observations are consistent with potential functional relevance of these loci and could inform future experimental studies to explore their roles in BMI plasticity.

Beyond genes, genetic interactions involving variants in promoter and enhancer regions may be associated with trait variability by modulating gene expression^23,24^. Two key findings in our study underscore this. First, several loci implicate lncRNAs: three BMI-associated loci map to LINC00880, and one, the top locus for infant length, is in suggestive interaction with a lncRNA variant in RPL34-DT. LncRNAs can act as cis-regulatory elements^25^ specifically during development^26^, with some functioning as enhancers^27^. Second, nearly all non-additive lead loci are noncoding variants enriched for intra-chromatin interactions with known enhancer and promoter regions. Together, these findings suggest a potential link through which noncoding variants may influence trait plasticity.

Epigenetic regulation is another mechanism that may contribute to developmental plasticity. DNA methylation is a major regulator of gene expression, with effects that vary over environmental exposures and time^28,29^. Approximately 34% of methylation QTLs are under genetic control, with the vast majority acting in cis^30^, similar to enhancers and promoters. Our empirical results align with these findings. This is especially evident for infant length, where the HIST1/2 histone gene clusters and canonical pathways related to positive and negative epigenetic regulation and histone acetylation and deacetylation, are enriched.

Across species, cis- and trans-regulatory elements are key drivers of gene expression variability and have been implicated in epistasis^23,31^. In *D. melanogaster*, cis-acting regulators are more prevalent and have stronger effects than trans-regulatory elements^23^, whereas the opposite pattern is observed in *S. cerevisiae*^31^. While our results demonstrate an abundance of cis-regulators, the inter-chromosomal interaction between MTMR11 and RPL34-DT indicates that trans-acting regulation may also contribute to non-additive genetic variation in humans.

We applied our custom-built models to a broad selection of traits, from anthropometric to cognitive. This allowed us to examine the degree to which the patterns of association are shared between these two types of traits, or if there are systematic differences between them.

One difference is the test statistics. There were more hits for anthropometric (driven largely by BMI) than cognitive traits, with anthropometric traits having cleaner association signals, overall. This likely reflects measurement error inherent in cognitive tests, even for psychometrically-sound instruments, such as the national tests in Norway. Another difference is the lack of external validation of putative plasticity loci for cognitive traits. Because additive loci for math and reading overlap with previous GWAS on EA and related traits, the lack of external validation likely reflects the limited number of vQTL and epistasis studies on cognitive and related traits, rather than a reflection of spurious association signal.

At a global level, non-additive, time-varying and additive genetic variation are intertwined, suggesting that additive genetic effects may partly capture interaction effects. This has important implications for the field. First, variance decomposition techniques used to estimate narrow-sense heritability assume genetic additivity. Our empirical results suggest that this may not always be the case. Second, although there are exceptions^32^, most Mendelian Randomization (MR) studies assume genetic variants only affect trait means. However, variants may have dynamic effects on traits; such that at certain timepoints they may affect trait levels, but at others also affect variability. This violates one of the key assumptions in MR studies (i.e. horizontal pleiotropy). And finally, studies using polygenic indices (PGIs) for interaction testing may be suffering from false negatives. The variants comprising PGIs are mostly derived from additive GWAS and may be missing signals from genomic loci that are involved in gene-environment and epistatic interactions. In light of our findings, greater caution is warranted in interpreting estimates from methods using variance decomposition and evaluating results from MR and PGI interaction studies.

There are several limitations in our study that should be noted. First, although our pipeline yields reliable and precise estimates, it is computationally demanding. Association testing for one phenotype across ∼65,000 individuals with three measures requires approximately 100,000 CPU hours. Second, our model currently fits only a linear trend over time, limiting its application to traits with non-linear trajectories. We plan to optimize our pipeline in future work and allow for non-linear trends to be modelled, especially in datasets with more time points. We provide our analytic code open source, which may serve as a template for the research community in the meantime. Third, our pipeline is calibrated to normally distributed traits. Using it for non-normal traits requires transformation. Scale transformations, however, may remove genuine genetic signal while failing to remove statistical artifacts that drive spurious correlations^6,4^. In ^6^, the authors implemented a simple solution that decorrelates additive from non-additive effects in BMI. This is, however, not fully tested and validated across scenarios and datasets. Given that we do not find evidence that the non-additive effects for log-transformed BMI in our study are driven by the general mean-variance relationship (**Supplemental Figure 11b**), and because additive effects may partly capture non-additive effects, we did not apply this additional correction. Finally, our results may lack generalizability, especially for loci that are not externally validated. This is particularly relevant for cognitive traits, where none of the putative plasticity loci overlapped with independent studies. Replication in an independent cohort with a sufficiently large sample and comparable test score measures is desirable.

A key strength of our study is the use of multiple approaches in addressing our main research questions. The concordance of association signals between our pipeline and trajGWAS lends internal validity to our findings, and the use of a simpler longitudinal model addresses some of the limitations discussed above. For instance, the validated BMI loci are robust to scale transformations since their effects do not rely on trait variance. Importantly, the inferences we draw from empirical results across different approaches are less likely to be affected by minor violations of key modelling assumptions.

In sum, our study offers potential mechanisms that may underlie developmental plasticity, implicating noncoding variants with potential regulatory roles, while providing empirical evidence that challenge the assumption that GWAS effect estimates are purely additive. Our findings open new avenues for functional genomic studies aimed at uncovering the molecular mechanisms that drive complex trait plasticity.

## METHODS

### Simulation

#### Bias, power and false discovery rate

Using *N = 10,000* individuals and *t = 3 time points*, we evaluated our log linear variance model and trajGWAS^11^ under six data generating models, wherein a trait with and without heteroscedastic residual due to a locus or single nucleotide polymorphism (SNP) is influenced by: a random intercept (**scenario 1**), by a random intercept and random slope (**scenario 2**), and by random intercepts in the mean and residual variance (**scenario 3**). To achieve this, we used:

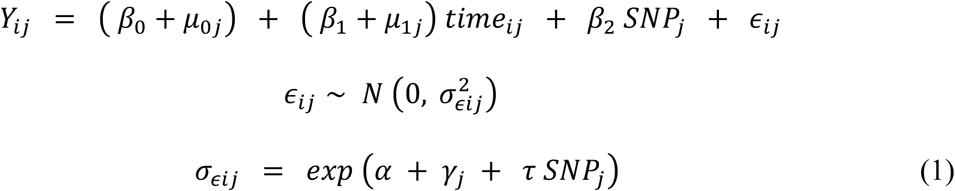

where *Y*, *I,* and *j* are as noted above, 𝛽_0_ is the intercept, 𝛽_1_ is the fixed effect of time on *Y*, 𝛽_2_ is the fixed effect of SNP on *Y,* and 𝜇_0j_ and 𝜇_1j_ are random terms in the mean. SNP was generated as noted above. The error term is parameterized as standard deviation denoted by 𝜎*_𝜖ij_*, which assumes a linear form on the log scale, and is comprised of a constant, 𝛼, a random intercept, 𝛾*_j_*, and a fixed effect, 𝜏, quantifying the heteroscedasticity due to SNP. The random terms are

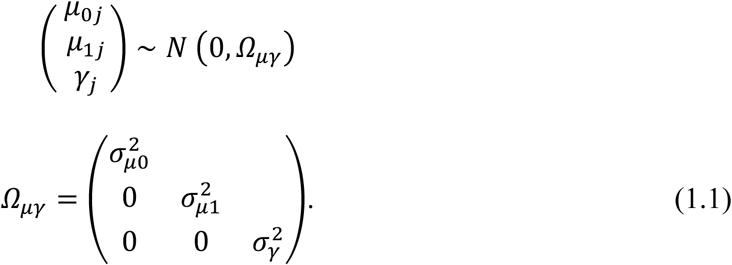

where the diagonals are variance terms (for the random intercept, 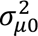, random slope, 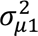, and random intercept in residual variance, 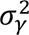) and the off diagonals are covariance terms, all set to zero.

To generate correlated phenotypes, we set 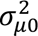 *= 0.8,* 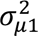 *= 0,* 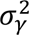 *= 0* for **scenario 1**; 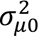 *= 0.8,* 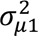 *= 0.04,* 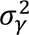 *= 0* for **scenario 2**; and 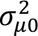 *= 0.8,* 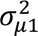 *= 0,* 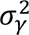 *= 0.04* for **scenario 3**. In all scenarios, all other terms were set to 0, except for 𝜏 ∈ *{0, 0.01, 0.05, 0.1*}*. Only the starred item was used for bias analysis. Each data generating model was executed 100x and the resulting datasets were saved and analyzed, first, using our log-linear variance model, then using trajGWAS. In all tests, a random intercept in the mean was fitted by our model, whereas random intercepts in the mean and the variance were fitted by trajGWAS. Bias and power were evaluated following ^33^. Power curves were generated using ggplot2.

### Ethics

Data from the Norwegian Mother, Father and Child Cohort Study (MoBa) and Statistics Norway (SSB) were used via the project SUBPU, which is approved by the Regional Committees for Medical and Health Research Ethics (ref. 2017/2205). The University of Oslo is responsible for the data handling in SUBPU and has conducted a Data Protection Impact Assessment (DPIA) in collaboration with the Norwegian Agency for Shared Services in Education and Research (Sikt; ref. 962088). The data access and management of SUBPU is financed by the Research Council of Norway, the European Research Council, and the Department of Psychology at the University of Oslo.

### The Norwegian Mother, Father and Child Cohort Study (MoBa)

MoBa is a prospective population-based pregnancy cohort study conducted by the Norwegian Institute of Public Health^34^. Participants were recruited from all over Norway from 1999 to 2008. The women consented to participation in 41% of the pregnancies. The cohort includes approximately 114,500 children, 95,200 mothers and 75,200 fathers, of whom 76,577 children, 77,634 mothers, and 53,358 fathers were genotyped and passed QC. MoBa is regulated by the Norwegian Health Registry Act.

#### Quality Assurance

Detailed individual and SNP quality control were discussed elsewhere^35^. In brief, duplicates, outliers in the subpopulation/ancestry PCA and those with discordant sex, heterozygosity ≥ 3 SD, and call rate < 98% were removed. Indels, strand ambiguous, duplicate SNPs were likewise removed along with SNPs with call rate < 95%, MAF < 1%, and HWE p-value < 1.00 × 10^−6^. Imputation was done in IMPUTE4 using HRC release 1.1. Only SNPs with INFO SCORE ≥ 0.8 were kept. Variants not found in the reference panel or with discrepant MAF > 0.2 relative to the reference panel were removed. Since MoBa contains complex interfamilial relations, additional checks excluded individuals with excessive cryptic relatedness and Mendelian error > 5%. Because MoBa genotyping spanned multiple projects across many years, SNPs were harmonized and those with plate, batch and imputation batch effects were excluded. A total of 6,981,748 hard called genotypes (threshold = 0.7) mapped to build GRCh37 (hg19) from 207,569 participants passed QC and were taken forward for association testing.

### Phenotypes

#### Early growth traits

Infant length and weight were extracted from MoBa questionnaires 4 and 5 completed by the mother when the child was 6 months and 12 months old, respectively. BMI was derived by dividing weight (kg) by infant length squared (m^2^). For infant length and BMI, outliers at each time point, defined as observations ± 3.5 SD or more from the sample average, were removed. Individuals with zero or decreasing length, and/or weight changes greater than 2 kg, were treated as implausible and excluded. Only individuals with observations from all timepoints were included. To minimize confounding of variance effects due to the general mean-variance relationship, BMI was log-transformed prior to analysis following ^4^ (**Supplemental Figure 3b**). For robustness, we checked if the minimal skew that persisted after transformation may have impacted our test statistics (**Supplemental Figure 11b**).

For the main analyses, we retrieved infant length and weight measures of infants aged 8, 12 and 18 months. The motivation for choosing these timepoints is 3-fold. First, these measurements were taken during routine post-natal checkup and recorded by a nurse in the health card. The mothers then used this card to complete the questionnaires, making these measurements less prone to error. Second, this timeframe contains the largest number of observations in our sample with almost the least amount of missingness (**Supplemental Table 1**). Third, the timeframe from 8 to 18 months captures a linear trajectory in BMI observed across multiple cohorts^18,36,37^. This allowed us to estimate heteroscedastic effects due to SNP using our model (**Equation 2**) while minimizing potential influences from nonlinear trends.

We found loci with time-varying effects from birth when annotating hits from our log linear variance model. Since birth length and BMI measures have the greatest number of observations, we modeled them together with growth trait measures from 8 and 12 months to maximize power and to capture a reasonably linear trend in developmental trajectory.

#### Cognitive traits

National test scores for math and reading were extracted from Statistics Norway (SSB) and linked to MoBa. Students with an exempt status were excluded from the analysis (∼5%). Exempt status is given to 1) students with special education needs and 2) non-Norwegian students who cannot participate in the tests since they are available only in Norwegian. The average was taken for students with more than 1 score recorded (i.e. likely repeaters) (∼0.1%).

The scores were standardized at the population level by school subject, grade level, the year the student took the exam and sex. Scores ± 3.5 SD from the average were considered outliers while non-exempt students who scored 0 were considered implausible and were removed. Participants with a missing test score were also excluded. Grade level was re-coded as time centered at grade 5.

Students in Norway are placed in a grade level based on age. Since 2007, around 95% of students at grades 5, 8 and 9 take a battery of tests at the beginning of the school year aimed at tracking and quantifying their skill in math, reading, and English independent of school curricula. The scores do not have a bearing on a student’s placement, thus represent the child’s aptitude and knowledge level in these subjects not attributable to motivation or test preparation skills. The tests are 40-60 items long. The test-retest reliability of the math and reading test across timepoints ranges between 0.7-0.85^38^. For a detailed discussion on the psychometric properties of these tests, we refer readers to ^39^.

Standardized math and reading scores are skewed which may drive genetic effects on residual variance. They were inverse normal transformed at a later step as a check of the robustness of association results (see below).

### Variant discovery

#### Log-linear variance model

A SNP involved in interactions has been shown in silico and empirically to induce an effect on both the level and variability of a trait^2–8^. Cao et al. ^5,40^ and Roennegård et al.^3,41,42^ have also shown that power is higher when SNP effects on trait level and variability are modeled simultaneously than separately. Given these, we implemented a heteroscedastic mixed effect model that jointly estimates a SNP’s effect on the mean (i.e. additive) and residual variance (i.e. non-additive) of a trait as follows:

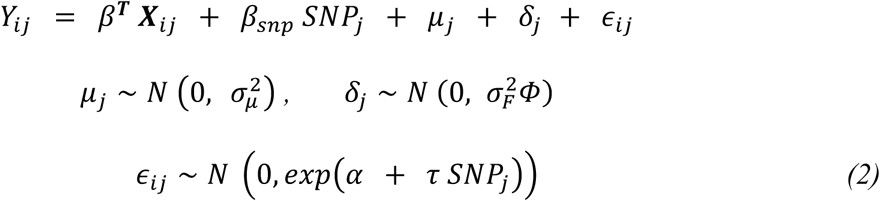

where *Y* is the trait of interest, *i* indexes time point, and *j* the person; ***X****_ij_* is an *n x k* matrix of covariates comprised of time°, year test was taken°*, sex°, batch, and 10 PCs, with their fixed effects denoted by the vector *𝛽^T^*; *𝜇_j_* is the random intercept representing an individual’s deviation from the sample mean, with 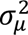 its effect on the trait level; 𝛿 is the genetic value, with variance 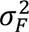 and Φ representing twice the kinship matrix of size *n x n*, and *𝜖_ij_* is the error term assumed normally distributed but allows for SNP heteroscedasticity to be estimated, here represented by 𝜏. The random terms are assumed independent, and 𝜏 is expressed as log-linear standard deviation along with the constant 𝛼.

For computational efficiency and to maximize N, **Equation 2** was implemented in 2 steps following ^5,40^. First, the phenotype for each time point was regressed on the GRM (constructed using ∼500,000 directly typed or imputed SNPs with INFO SCORE ≥ 0.975 and 0.01 ≤ MAF < 0.5) using the –reml and –reml-pred-rand option in GCTA^43^. This step was necessary to keep the influence of the dense and complex interfamilial relations in our dataset from artificially deflating the standard error of the estimated SNP effects (**Supplemental Figure 4**). This step also removed linear and nonlinear time trends, i.e. age and age^2^ (**Supplemental Figure 5, Supplemental Table 2**). In the second step, association testing was performed, where individuals with missing genotype at each locus were excluded. To detect SNPs affecting the level and/or variability of the trait, a joint likelihood ratio test was performed using the anova function, which compared a base model without SNP against the full model (**Equation 2)**.

° *Math and reading scores were standardized at the population level. The effects of year test was taken and sex were not completely removed* (**Supplemental Figure 2**)*, hence were added as covariates in the association testing step*.

* *Covariate used only for math and reading.*

#### Controls for population structure

To control for population stratification, we adjusted the p-value from the association testing described above by the genomic control factor^44^. Ko et al.^11^ have shown in simulation that uncontrolled population structure in the residual may drive p-value inflation, thus increasing the false positive discovery rate. In line with this, we took all significant SNPs and re-ran them through our pipeline, including 10 principal components (PCs) in the residual at this step. SNPs that survived these controls were taken forward for further testing.

#### Variants with non-additive effects

To ensure significant SNPs from the prior step are associated with trait variability and not just trait level, we performed an additional likelihood ratio test. Specifically, we compared a base model including only the SNP’s mean effect against the full model (**Equation 2**).

For math and reading, an additional robustness check was conducted. SNPs significantly associated with trait variability were re-tested for association using inverse normal transformed phenotypes.

#### Genotype-by-time model

Genetic effects on trait variability are more difficult to detect because they have larger sampling error than genetic effects on trait level^2^. Recently, longitudinal GWAS have found genetic variants with time-varying effects^36,45^. This may be due to epistatic or gene-environment interactions. We fitted a SNP-by-time model to our longitudinal phenotypes as follows:

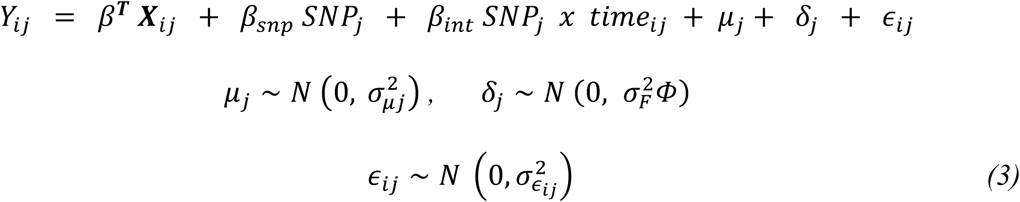

where 𝛽*_int_* is the effect of SNP-by-time interaction. Since our main interest is on the parameter 𝛽*_int_*, genotypes were centered and standardized by the MAF. All other variables are as noted in **Equation 2**, except the residual variance, 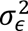, assumes homoscedasticity. Note that *X_ij_* includes time as covariate.

### TrajGWAS

Traits residualized on the GRM were imported to Julia and tested for association using the default options in trajGWAS (wald = FALSE, spa = TRUE). We fitted the model:

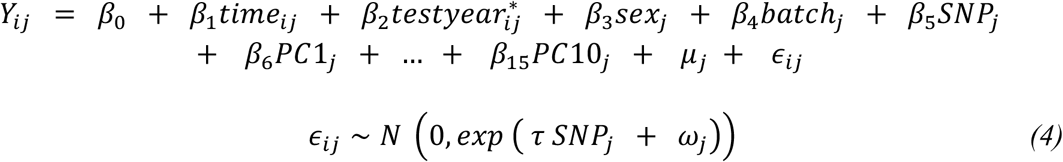

where *i* indexes time point, *j* individual and the 𝛽, coefficients represent the intercept and the fixed effect of each of the variable on the mean of the phenotype, *Y_ij_*. Both *𝜇_j_* and *𝜔_j_* are random intercept for the individual assumed i.i.d. with mean zero and covariance

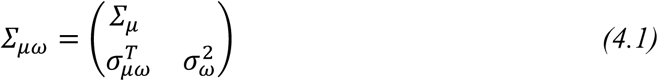

For each phenotype, we extracted the resulting Wald test p-value for comparison with the joint effect likelihood ratio test p-value from our variance model.

*\* Covariate used only for math and reading*.

### Biological annotation

We performed annotation in FUMA and MAGMA with genome build hg19 using the European reference panel in 1000 Genomes Project - Phase 3. Loci in the major histocompatibility region (MHC) were excluded. Lead loci were defined using a merge distance of 250 kb and an LD threshold of r² ≥ 0.6. Independent loci were defined using r² < 0.1. All gene types mapped to Ensembl v102 were included. All available datasets were used for eQTL and chromatin interaction (Hi-C) mapping, using a false discovery rate of 1 x 10^-6^. The complete list of Hi-C datasets is available in **Supplemental Table 15**.

### Correlation of additive, non-additive, and time-varying additive effects

We retrieved the effect estimates from our log-linear variance model as well as genotype-by-time model for each trait. Both additive and non-additive effects from our variance model were centered and standardized by the MAF prior to calculating pairwise correlations using cor.test().

In population vQTL studies, a SNP affecting trait variability may be driven by a neighboring SNP that affects the trait level if they have discordant MAFs – a phenomenon termed “phantom vQTL”^40^. To ensure that the correlation coefficients are not affected by LD, we extracted all SNPs in linkage disequilibrium with our non-additive SNPs using the LD option in PLINK (--ld-window 99999 --ld-window-r2 0.01). We used the mean effects estimated in our log-linear variance model to retrieve SNPs with suggestive genome-wide significant effect on trait level. We then compared this list against the list of SNPs in LD with non-additive SNPs.

### IT Infrastructure

Handling and processing of individual-level data were conducted on TSD (Tjenester for Sensitive Data), owned by the University of Oslo, operated and developed by the TSD service group - IT Department (USIT). All statistical analyses were carried out using Colossus provided by Sigma2, the National Infrastructure for High-Performance Computing and Data Storage in Norway.

### Software

Unless otherwise stated, simulations, data analyses, and plots were done in base R. Manhattan plots were generated using ggplot2 and the accompanying QQ plots using the *qqman* package. Circos plot was created using *BioCircos*. The package *bigsnpr* was used in handling genetic data and *nlme* in implementing our longitudinal variance model.

### URLs

Analytic pipeline: https://github.com/RPorneso/longitudinal_variance_GWAS

Pre-registration: https://osf.io/43mtv/overview (note inclusion of growth traits in May 2024) FUMA v1.5.2 (MAGMA v1.08): https://fuma.ctglab.nl

Summary statistics are available upon request.

## ACKNOWLEDGEMENTS

This work was supported by the European Union’s (EU’s) Horizon Europe research and innovation program under the Marie Skłodowska-Curie grant agreement (no. ESSGN 101073237), the EU grant agreement (grant no. 101045526, project GeoGen), and by Sigma2 (grant no. NS9867S). The Norwegian (336078, 288083, 331640) and Swedish Research Councils (2024-06499_VR) supported this work. We thank Michel Nivard for the fruitful discussions in the earlier phase of the project, Perline Demange for reading the pre-registration, and Clara Timpe and Oda van Jole for managing MoBa data. For generating high-quality genomic data, we thank the Norwegian Institute of Public Health (NIPH), the HARVEST collaboration, the NORMENT Centre at the University of Oslo, the Center for Diabetes Research at the University of Bergen, deCODE Genetics, the Research Council of Norway, the South-Eastern and Western Norway Regional Health Authorities, the ERC AdG, Stiftelsen KG Jebsen, the Trond Mohn Foundation, and the Novo Nordisk Foundation. We are grateful to all the participating families in Norway who take part in this on-going cohort study.

## AUTHOR CONTRIBUTIONS

R.P., E.Y., and A.H. conceived the study. E.E. provided technical expertise for model development and simulations. R.P. developed the analytical pipeline, performed simulations, conducted genome-wide analyses, ran functional annotations, interpreted results, and drafted the manuscript. All code/simulations independently verified by E.E. A.H. and E.Y. managed MoBa data access and governance. A.H., E.E., and E.Y. provided manuscript feedback. E.E. and E.Y. supervised the work.

