## Supplemental Figures for "Longitudinal modelling reveals widespread non-additive genetic effects underlying developmental plasticity"

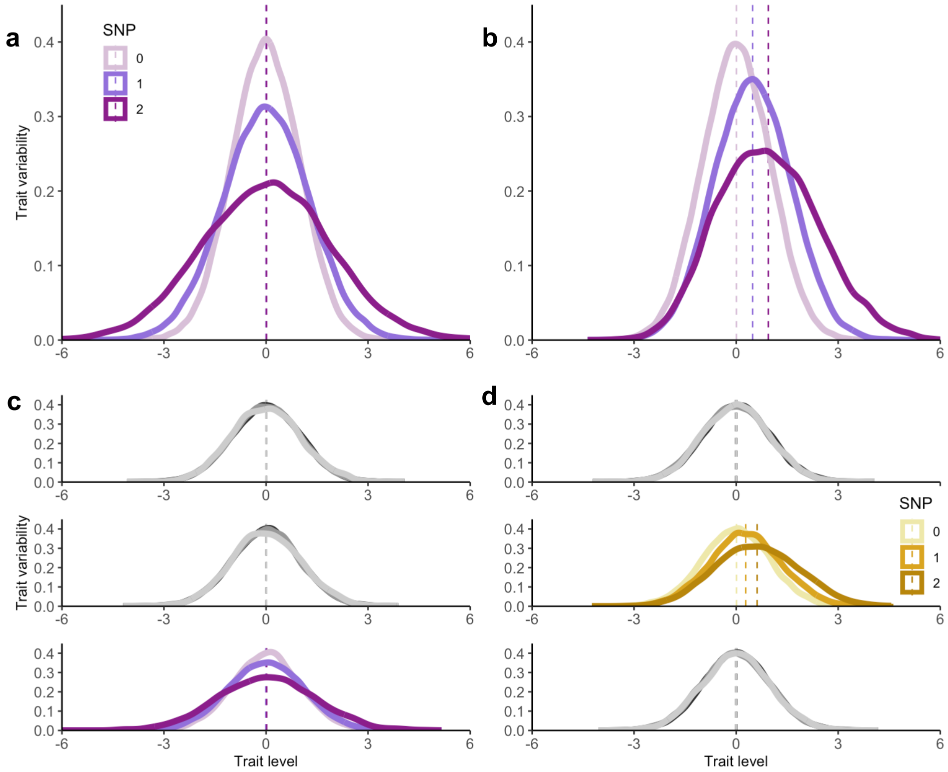


**Supplemental Figure 1. Effects of genetic interactions on trait level and variability.** A non-additive genetic effect (purple, gold) is simulated via gene–environment interaction (a, c) or epistasis (b, d). In (a) and (b), the interactions affect all time points, while in (c) and (d), the interaction effects are present only at a single time point. Dashed lines represent trait mean. In (a), gene-environment interaction influences only trait variability, reflected in the scale of the distribution, whereas in (b), epistasis also impacts trait level, which shifts the location of the distribution. Under the specific data generating models laid out in Supplementary Note 1, panels (c) and (d) demonstrate that epistasis, but not gene-environment interaction, induces a time-varying additive effect (gold) detectable in longitudinal GWAS that assume homoscedasticity.

**Supplementary Note 1: Gene-environment and epistatic interaction data generating models**

A locus or single nucleotide polymorphism (SNP) interacting with either the environment or another SNP generates heteroscedasticity[2–8]. To validate this, we simulated a genetic variant, *SNP ~ Bin(2, 0.3)*, and an environmental factor, *E ~ N(0, 1)*, for 60,000 individuals. We then generated a phenotype, $Y = \beta\ SNP\ x\ E + \epsilon$, where $\epsilon \sim N(0, \sigma^2)$, and $\beta = 0.8$. We tested for heteroscedasticity due to SNP using Levene's test with the Brown-Forsythe option from the car package. And we checked for the effect of SNP on trait level (mean) by running a univariate linear regression using the lm() function. For comparison, we simulated a phenotype under an additive genetic effect framework using $Y = \beta\ SNP + \epsilon$.

Since our main interest is in detecting genetic effects in within-individual variability, we repeated the above approach, simulating a phenotype with correlated measures (timepoints, *t = 3*):

$$Y_{ij} = \beta\ SNP_j\ \ x\ \ E_{ij} + \epsilon_{ij}$$

$$\epsilon_{ij} \sim N\left(0,\ \begin{matrix} 1 & \rho & \rho \\ \rho & 1 & \rho \\ \rho & \rho & 1 \end{matrix}\right).$$

Here, $i$ indexes time point, $j$ the individual, and $\rho$ (*0.8*) is the correlation between repeated measures. We ran a linear mixed model to verify that the random intercept captured between-individual variability.

We additionally tested if the above data generating models would induce time-varying genetic effects. We fitted a genotype-by-time model with correlated measures, testing scenarios where the interaction was present at all timepoints versus only a subset. We used the following data generating models for the latter case:

$$Y_{ij} = \beta^T\ SNP1_j\ x\ \ E_{ij} + \epsilon_{ij}$$

$$Y_{ij} = \beta^T\ SNP1_j\ x\ \ SNP2_j + \epsilon_{ij}$$

where $\beta^T = (0, 0.5, 0)$ or $\beta^T = (0, 0, 0.5)$ indicating that the genetic interaction is only present at time point 2 or 3. In all tests, we fitted main effects for SNPs and environmental factor along with their interaction.

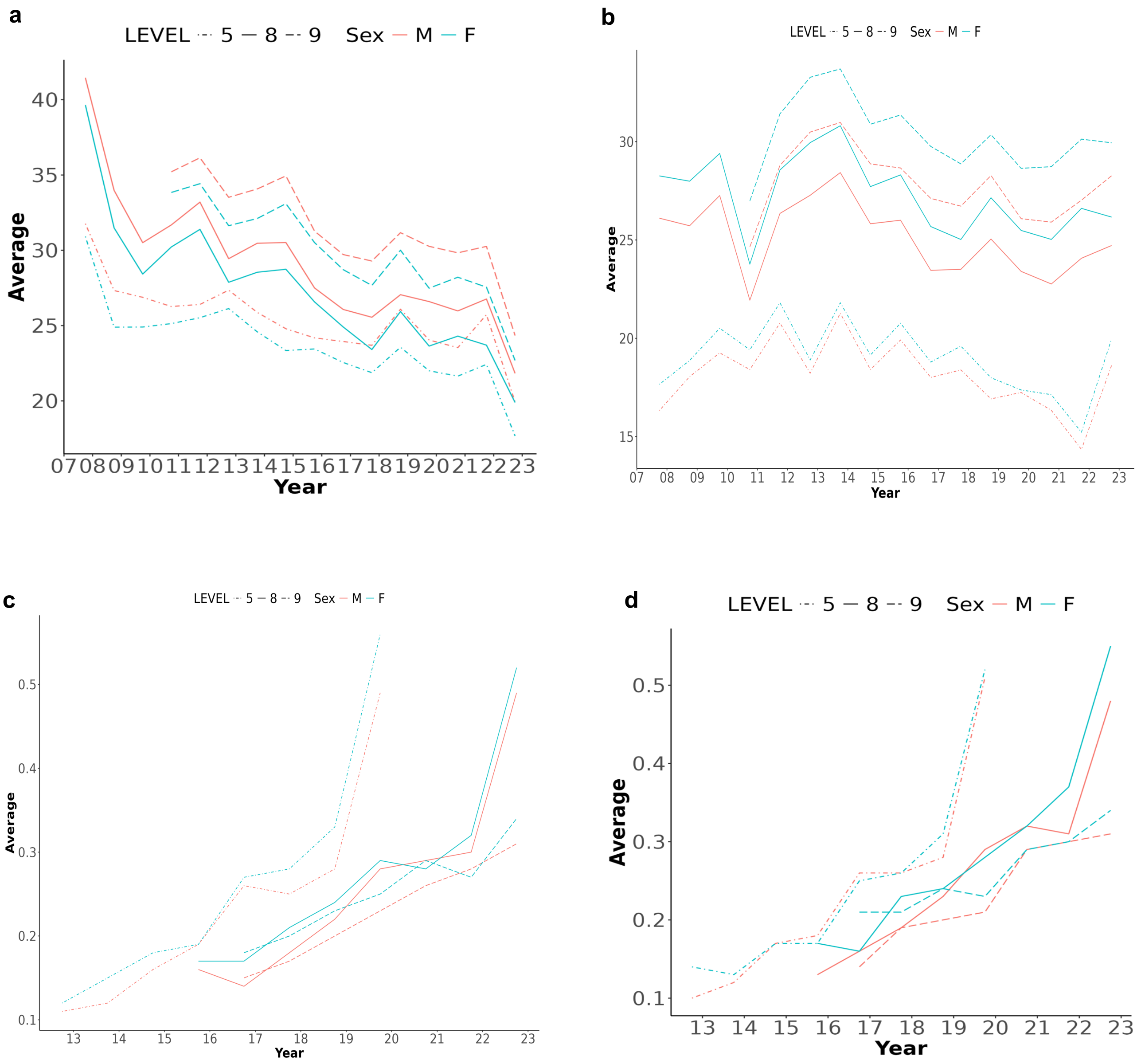


**Supplemental Figure 2. Yearly trend of math (a, c) and reading scores (b, d)**. In (a) and (b), raw scores from the entire population are shown. In (c) and (d), standardized scores are shown for the sample used in the study. Standardization at the population level did not remove the effects of year test was taken, grade level (i.e. age), and sex, reflecting the selection bias in the sample. Association testing included these as covariates to account for their effects accordingly.

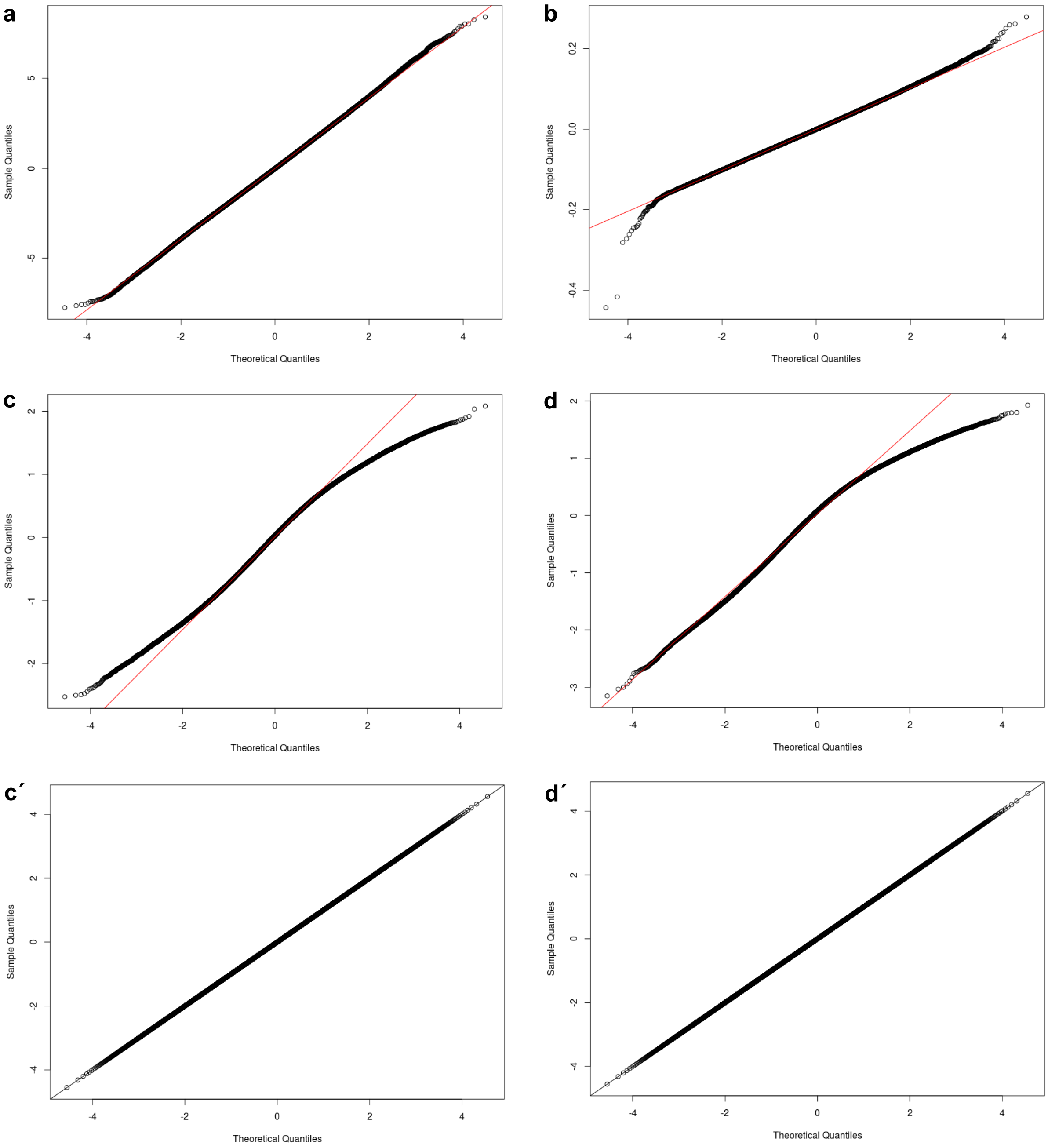


**Supplemental Figure 3. Trait distribution.** Normal QQ plot for residualized traits: infant length (a), log-transformed BMI (b), standardized math (c) and standardized reading scores (d). Both math and reading have negative skew, hence were inverse-normal transformed (c´, d´). Loci with significant effects on trait variability were re-tested for association after inverse normal transformation.

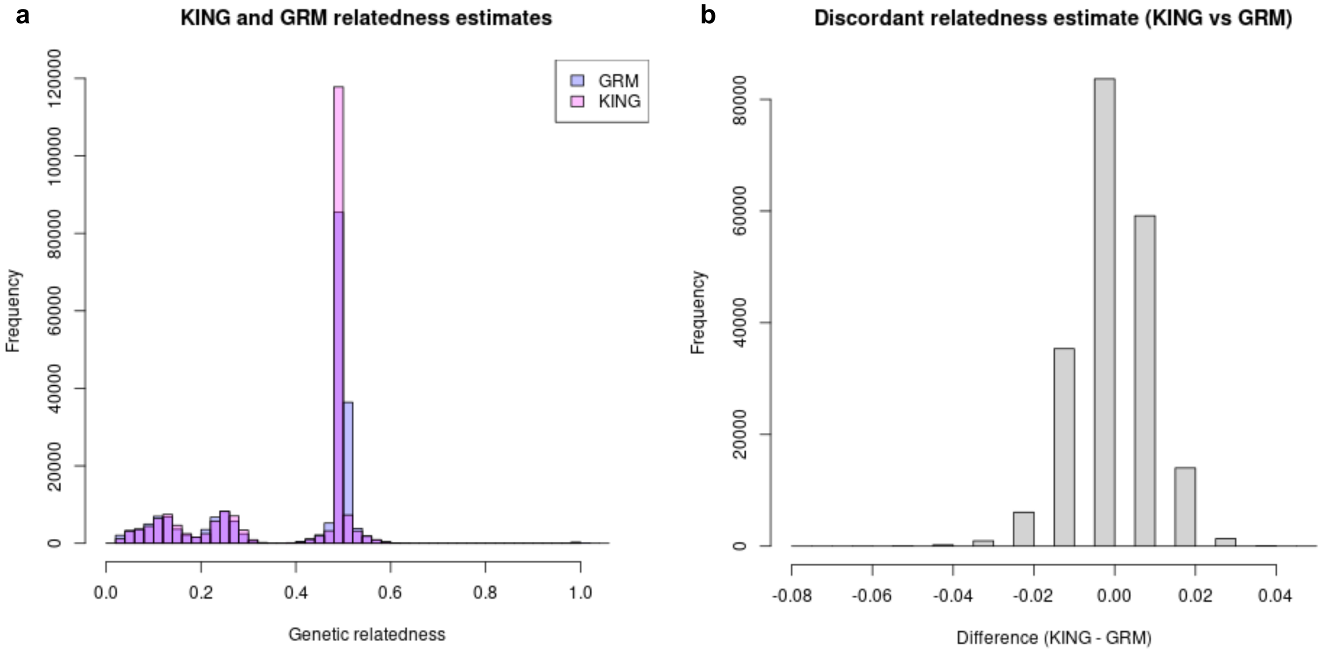


**Supplemental Figure 4. Sample relatedness using GRM versus KING estimates.** The dense interfamilial relations in our sample is visualized in (a), where a lower bound cut off of 0.03 was used. The relatedness estimates are largely the same between GRM and KING (b). Note that the GRM found more pairwise relations than KING (data not shown). Trait values were residualized on the GRM for each time point (age) prior to association testing. This step is necessary to make the observations independent from each other within each time point. The observations within individuals (i.e. across time points) are not independent, hence a random intercept was fitted at the association testing step.

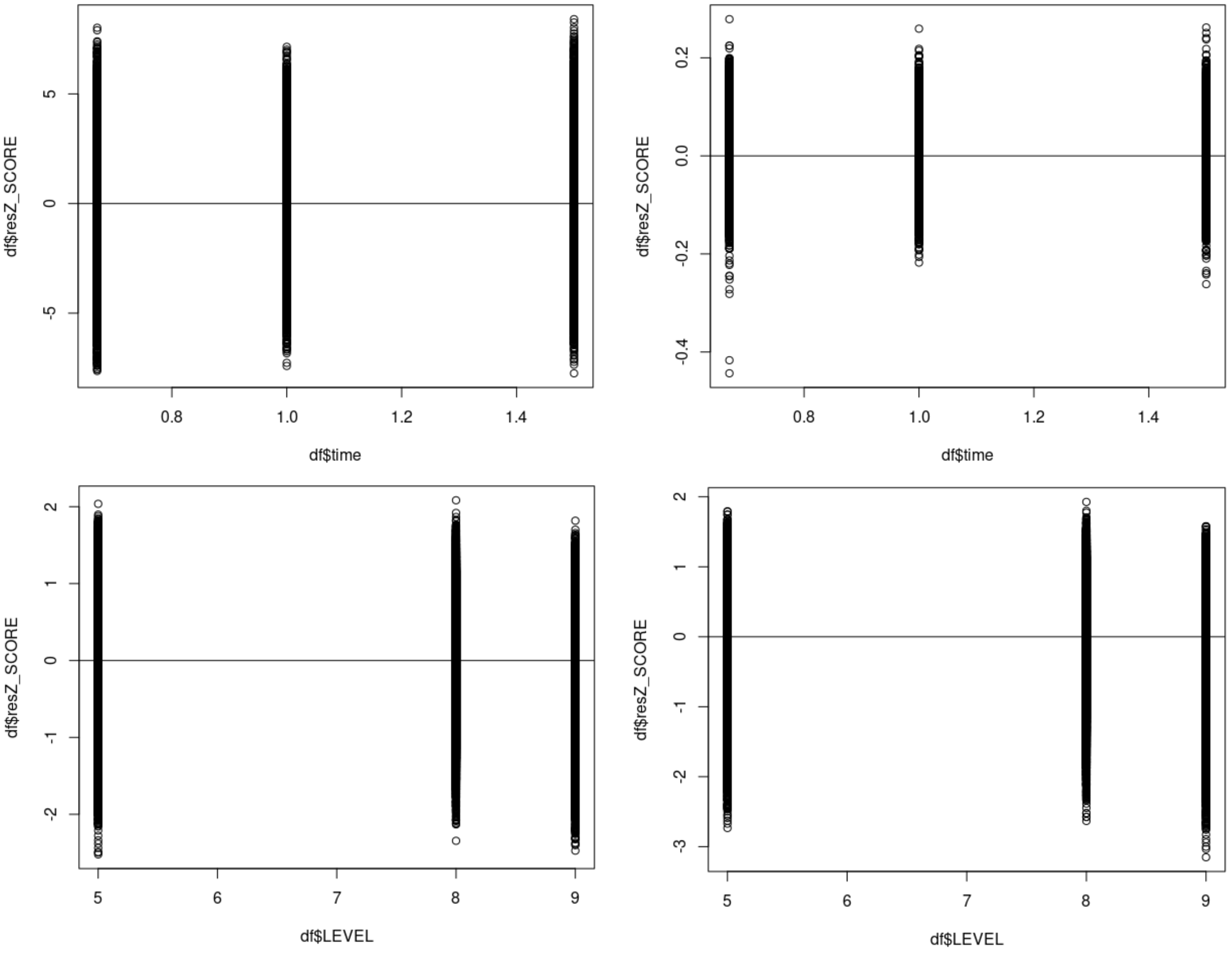


**Supplemental Figure 5. Age trend across traits.** Regressing the phenotypes on the GRM at each time point removed both linear and nonlinear time (age) trends: infant length (a), log-transformed BMI (b), standardized math (c) and reading scores (d). In (a, b), age is labeled time, whereas in (c, d) LEVEL corresponds to grade level. Norwegian students are assigned to a grade level based on age **(Supplemental Table 1).** Estimates from the mixed effect regression including the p-values, AIC, BIC and Loglik are provided in **Supplemental Table 2**.

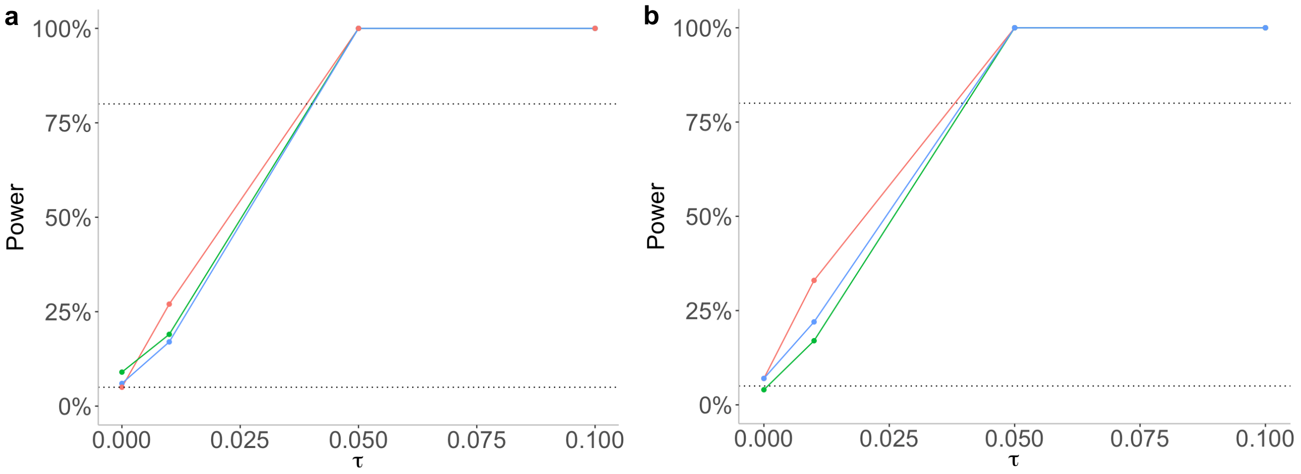


**Supplemental Figure 6. Log-linear variance model versus trajGWAS power and false positive rate under three data generating models (N = 10,000 and MAF = 0.3 ).** Log-linear variance model (a) and trajGWAS (b). Data generating models are color coded: red = random intercept, blue = random intercept and slope, green = random intercept in the mean and residual. The effect sizes (i.e. tau) are in standard deviation.

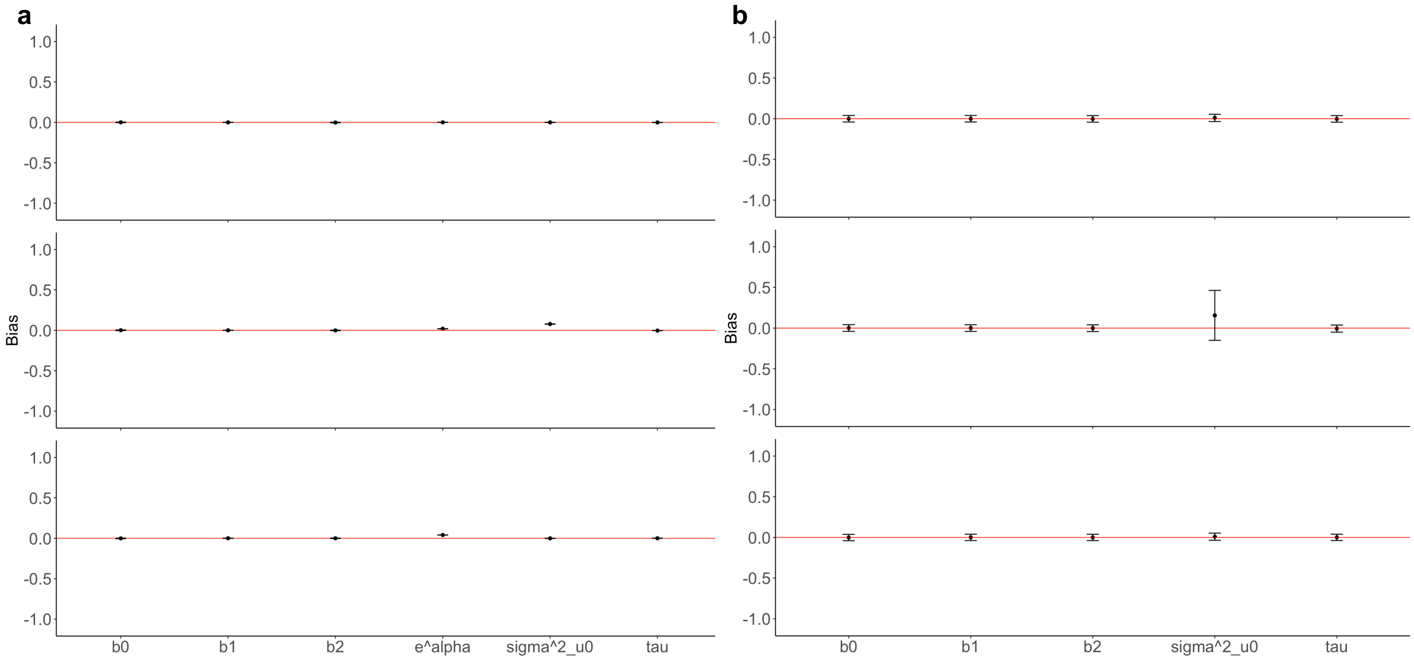


**Supplemental Figure 7. Log-linear variance model versus trajGWAS bias under three data generating models (N = 10,000 and MAF = 0.3).** Log-linear variance model (a) and trajGWAS (b). Tau is the estimated non-additive genetic effect. Not accounting for random slope and random intercept in the residual does not impact non-additive effect estimates. Unmodelled random effects are instead absorbed by the random intercept on the mean or by the residual variance (a). Note: TrajGWAS does not return residual variance estimates, i.e. e^alpha. Top = random intercept only, middle = random intercept and slope, bottom = random intercept in the mean and residual.

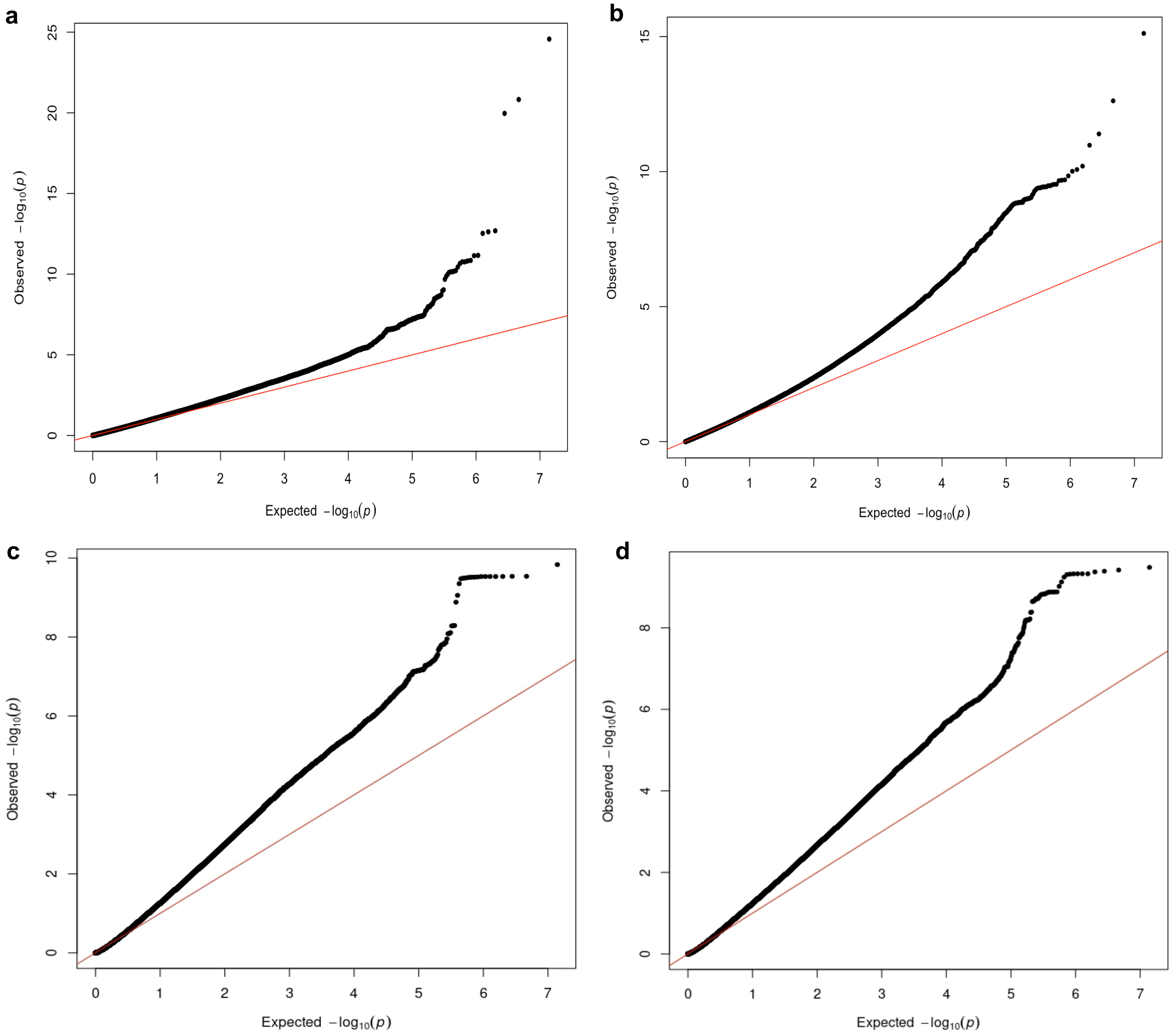


**Supplemental Figure 8. QQ plots after genomic control.** Infant length (a), BMI (b), math (c), and reading (d). P-values from the mean and variance likelihood ratio test are plotted on the y-axis. $\lambda_{GC} = 1$ across traits. These plots accompany Figure 2 in the main text.

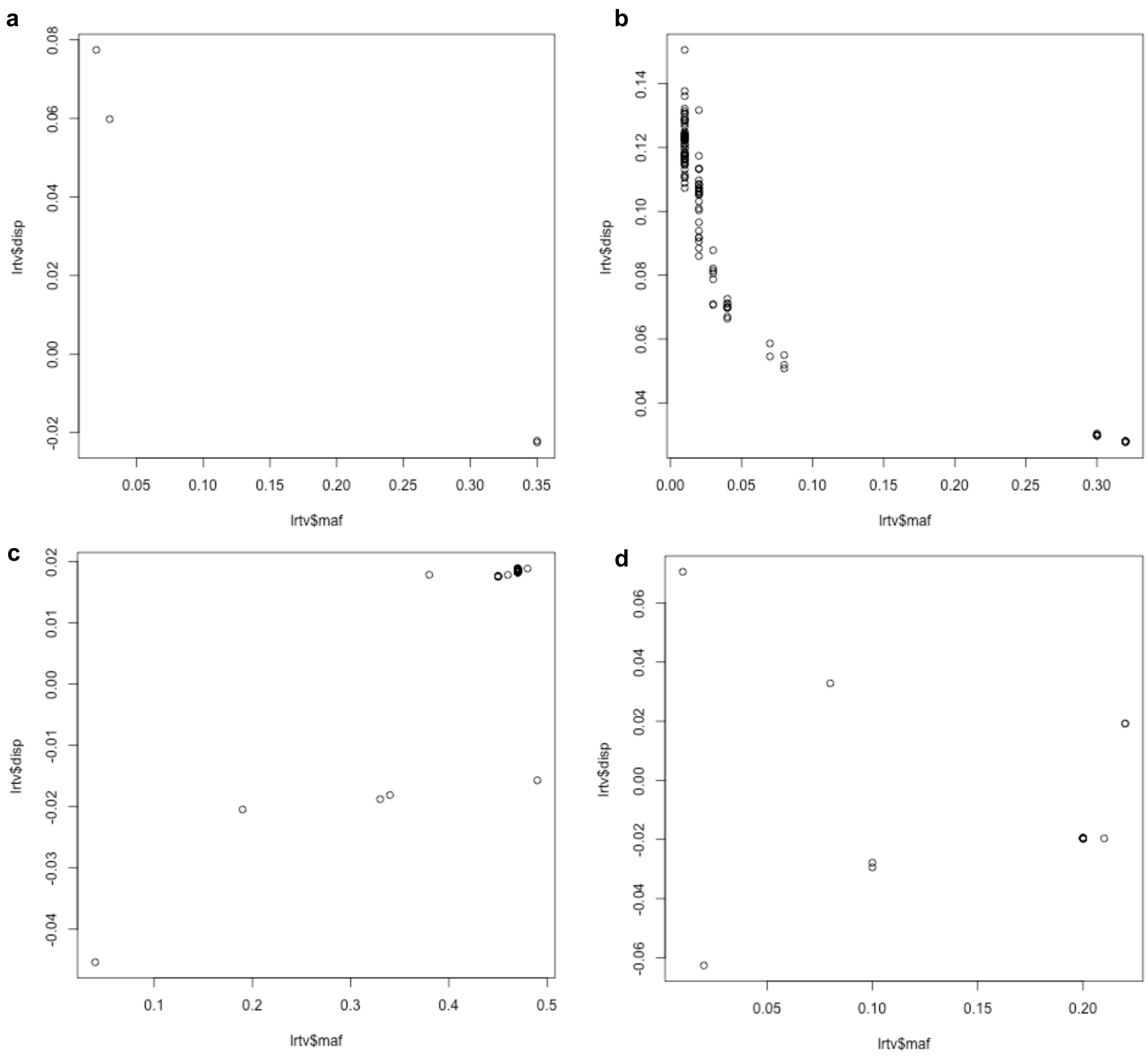


**Supplemental Figure 9. Non-additive effects by MAF.** Putative plasticity loci for infant length (a), BMI (b), math (c), reading (d). These plots accompany Figure 2 in the main text. Non-additive effect estimates (i.e. disp) in y-axis are in standard deviation.

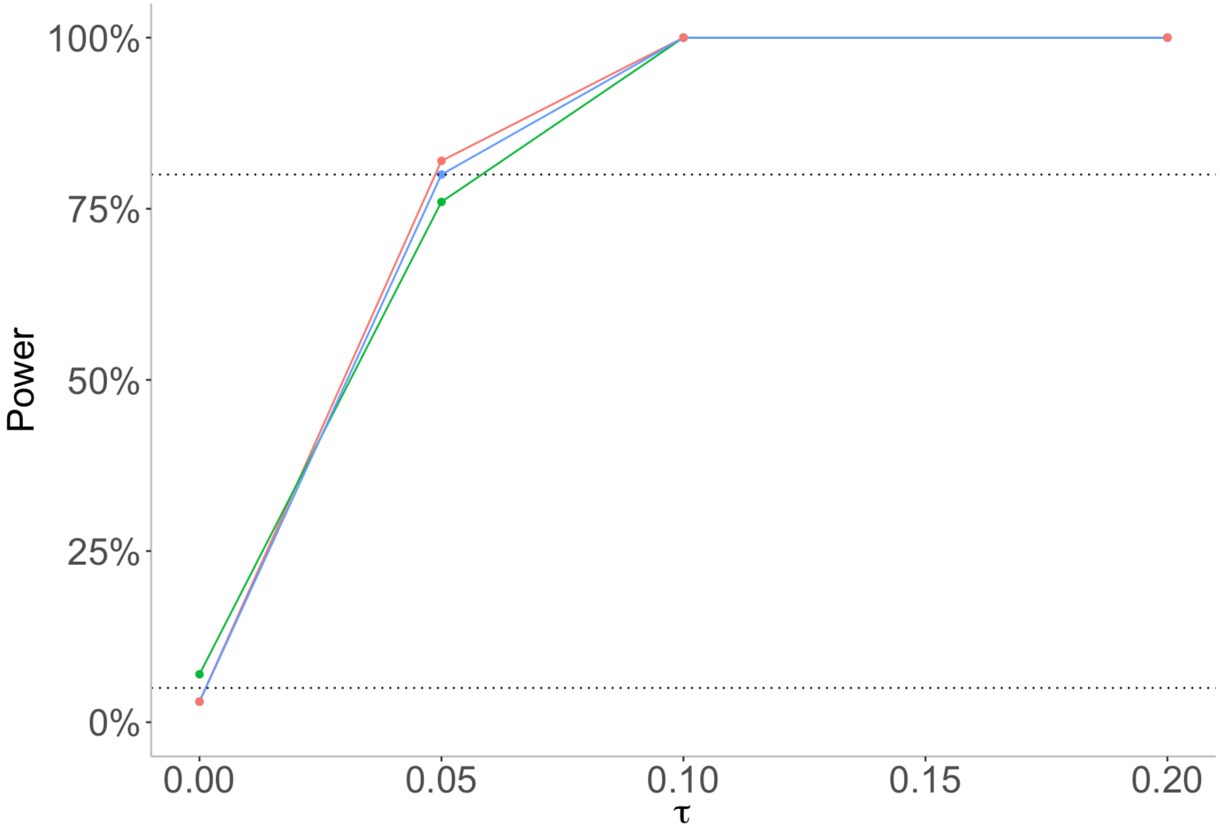


**Supplemental Figure 10. Log-linear variance model power and false positive rate under 3 data generating models (N = 50,000 and MAF = 0.01).** Data generating models: red = random intercept, blue = random intercept and slope, green = random intercept in the mean and residual. Non-additive effect sizes (i.e. tau) are in standard deviation.

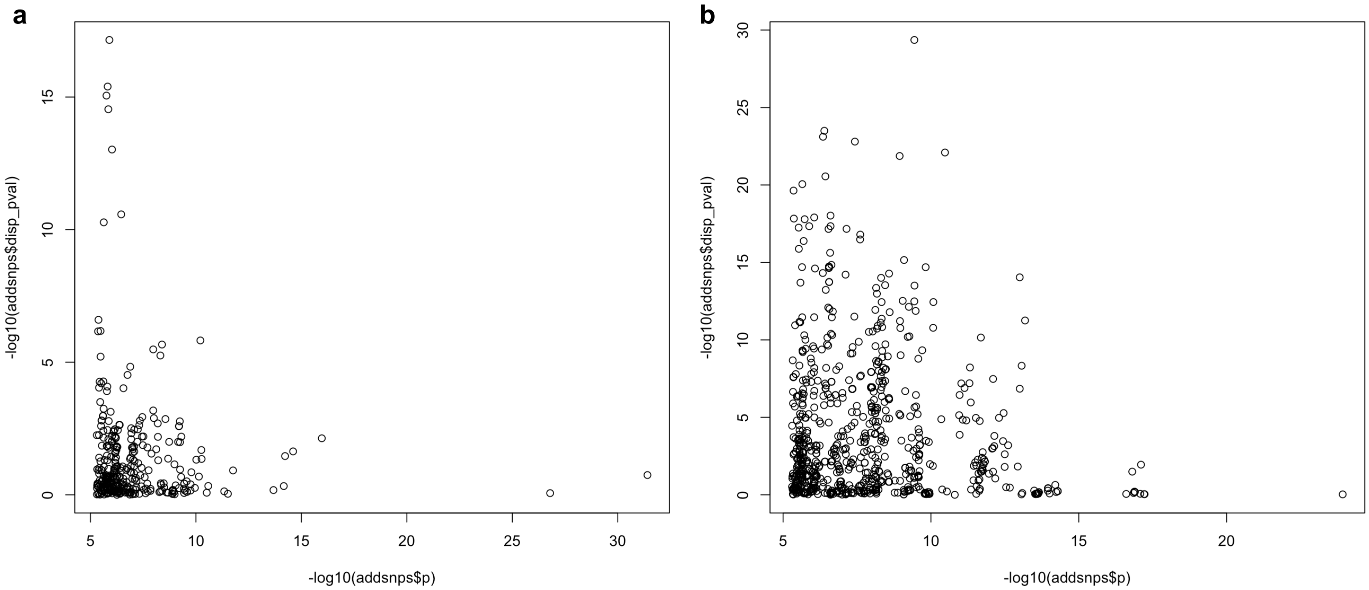


**Supplemental Figure 11. Non-additive genetic effects for growth traits are not due to general mean-variance relationship.** For non-normal traits, genetic effects on trait variability may be due to the general mean-variance relationship. The p-values of loci with significant additive effects are plotted against their non-additive effect p-values: infant length (-0.087, $p = 0.10$) (a) and log-transformed BMI (-0.138, $p = 8.33 \times 10^{-5}$) (b). The negative correlation in (b) suggests that the putative plasticity loci we discovered are not due to scale effects.

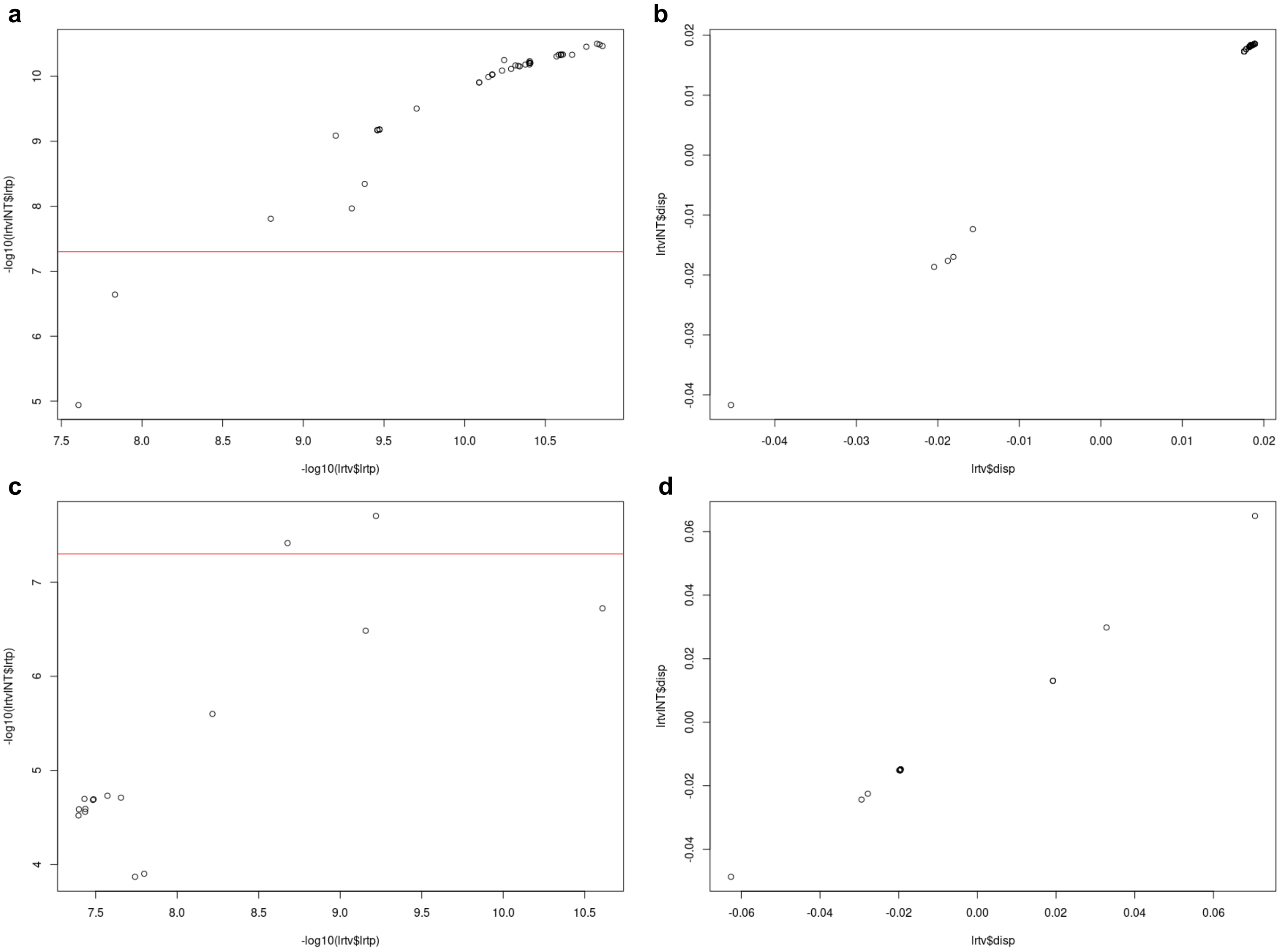


**Supplemental Figure 12. Putative plasticity loci for cognitive traits before and after inverse normal transformation.** Math (a, b) and reading (c, d). P-value before and after transformation (a, c). Red horizontal line corresponds to the genome-wide significance threshold. Effect sizes shown in (b, d), where the correlation coefficients are unity ($p < 2 \times 10^{-16}$): 0.999 (b) and 0.997 (d). The average decrease in effect size after transformation is 0.005 for both traits. Note the effects (i.e. disp) are in standard deviation with respect to genotype dosage = 0.

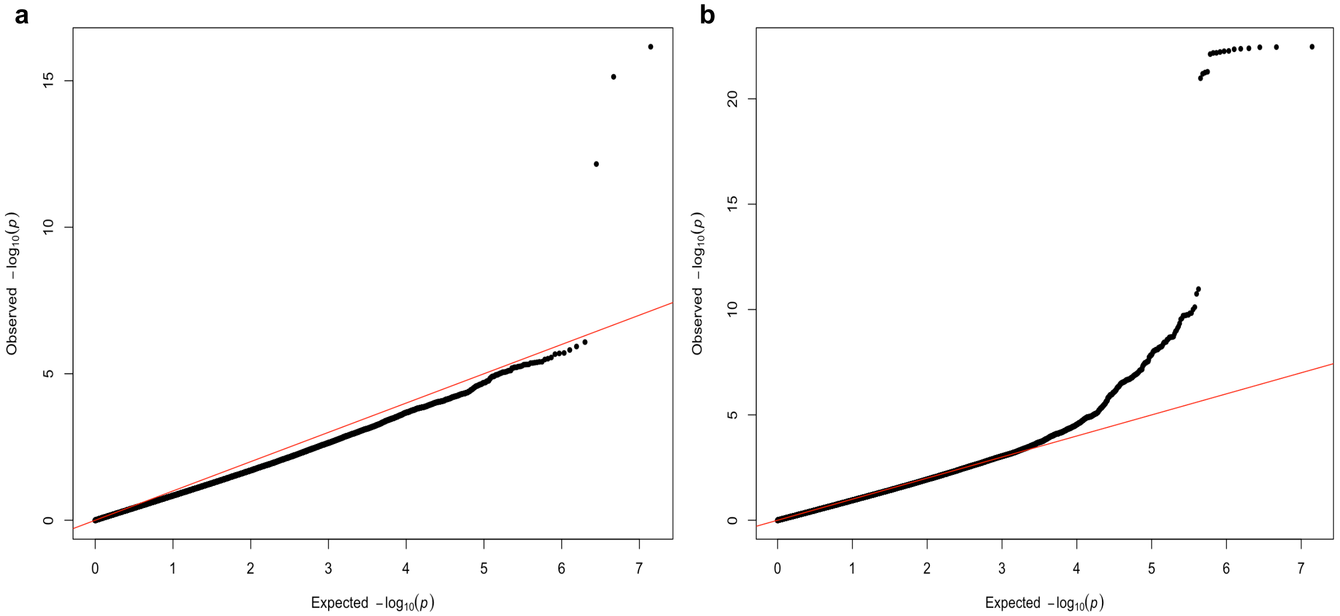


**Supplemental Figure 13. QQ plots (time-varying effects) for growth traits using measures from birth as baseline.** The p-values plotted in the y-axis are from the genotype-by-time parameter. There is a slight deflation in p-values. The traits were residualized on the full GRM prior to association testing. Leave-one-chromosome-out (LOCO) was not performed for computational efficiency and to account for the dense interfamilial relations in the sample. Note that genetic principal components (PCs) were included in the association testing step to control for population structure and other environmental confounding. Infant length (a) and BMI (b). These plots accompany Figure 5 in the main text.

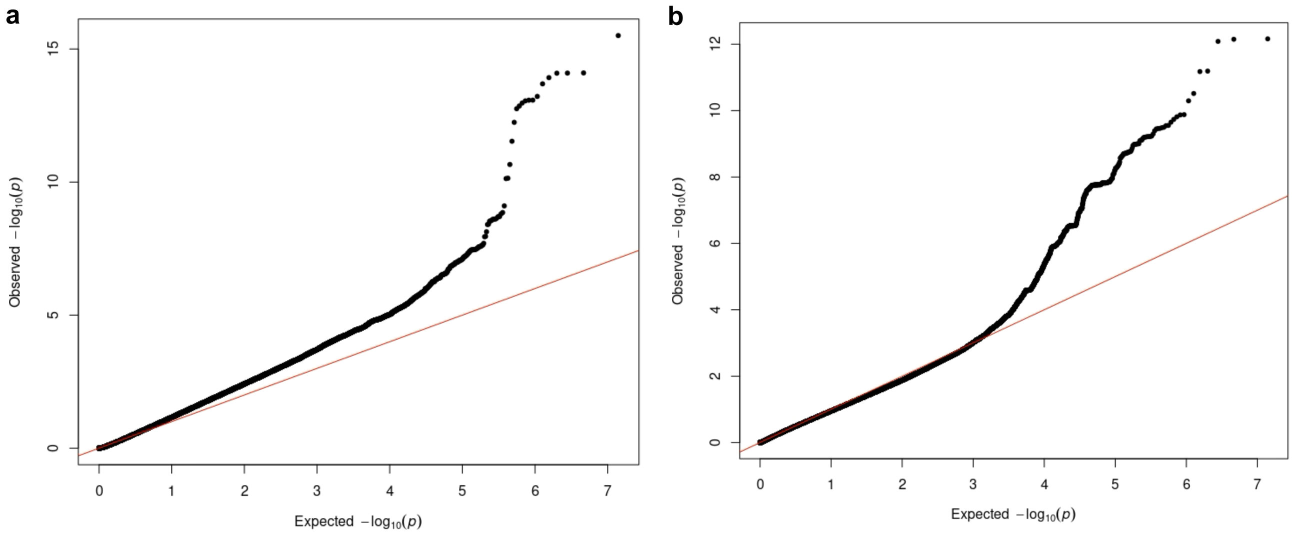


**Supplemental Figure 14. QQ plots after genomic control for growth traits using measures from birth as baseline.** Infant length (a) and BMI (b). P-values from the mean and variance likelihood ratio test are plotted on the y-axis.

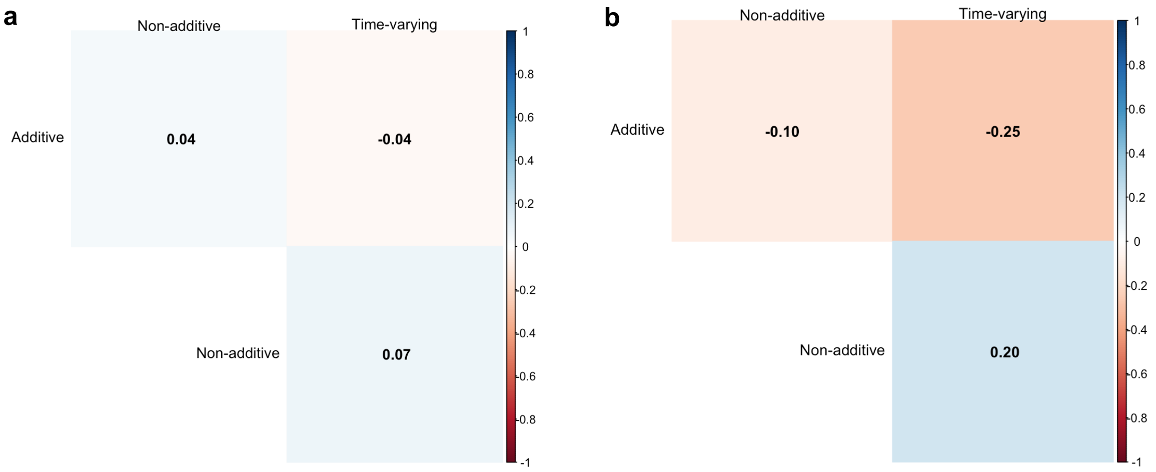


**Supplemental Figure 15. Correlation matrix for growth traits using measures from birth as baseline.** Infant length (a) and BMI (b). The sign of the correlation coefficients show a different pattern based on time points used.

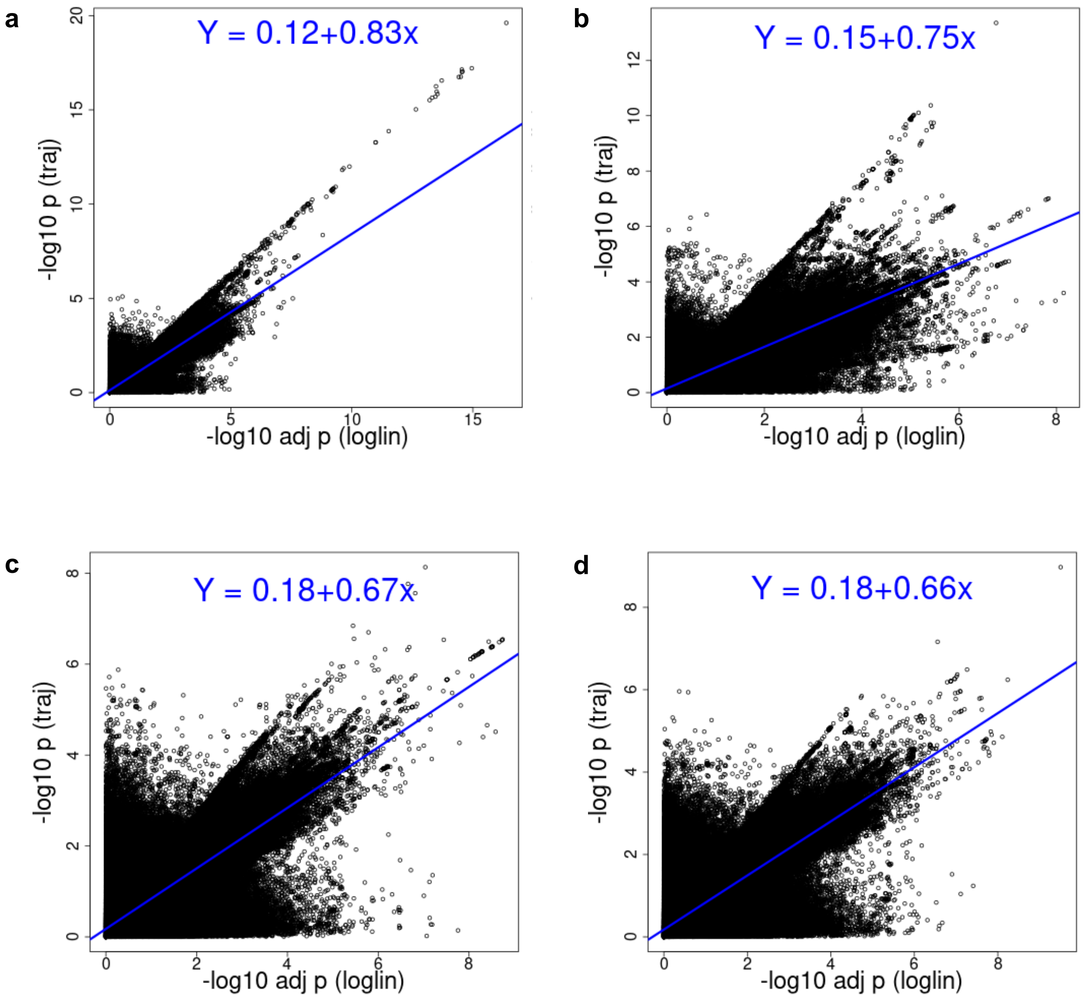


**Supplemental Figure 16. Comparison with trajGWAS.** P-value correlation of Wald test p-value from trajGWAS on the y-axis and LRT p-value after genomic control from our log-linear variance model on the x-axis. Infant length (a), BMI (b), math (c), and reading (d). The correlations in (a, b) are based on growth trait measures from birth, 8, and 12 months. All correlation p-values < 2e-16.

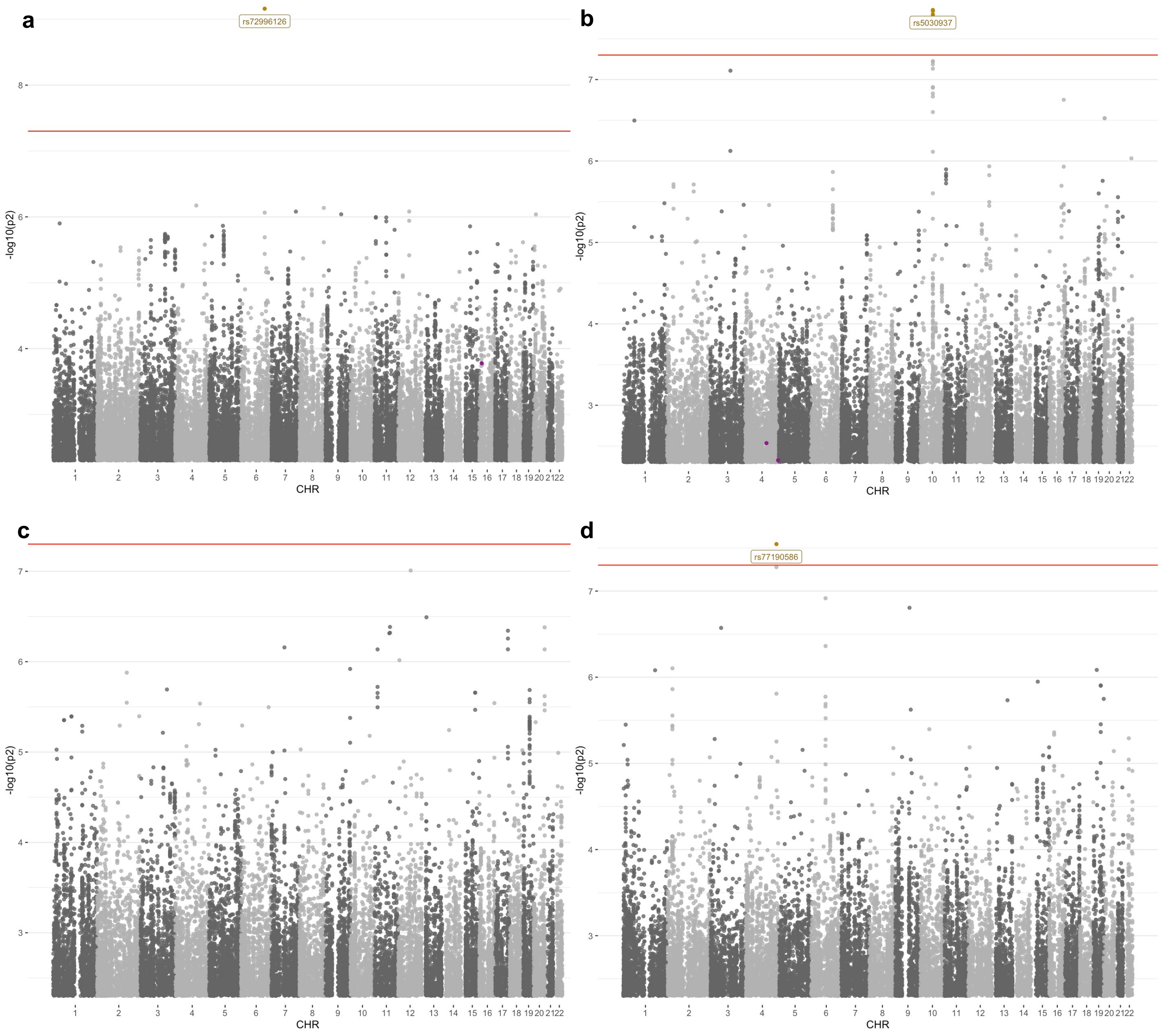


**Supplemental Figure 17. Loci with time-varying effects.** Genotype-by-time Manhattan plots for infant length (a), BMI (b), math (c), and reading (d). In (a) and (b), measures were taken at 6 , 8, and 12 months. QQ plots are available in Supplemental Figure 18.

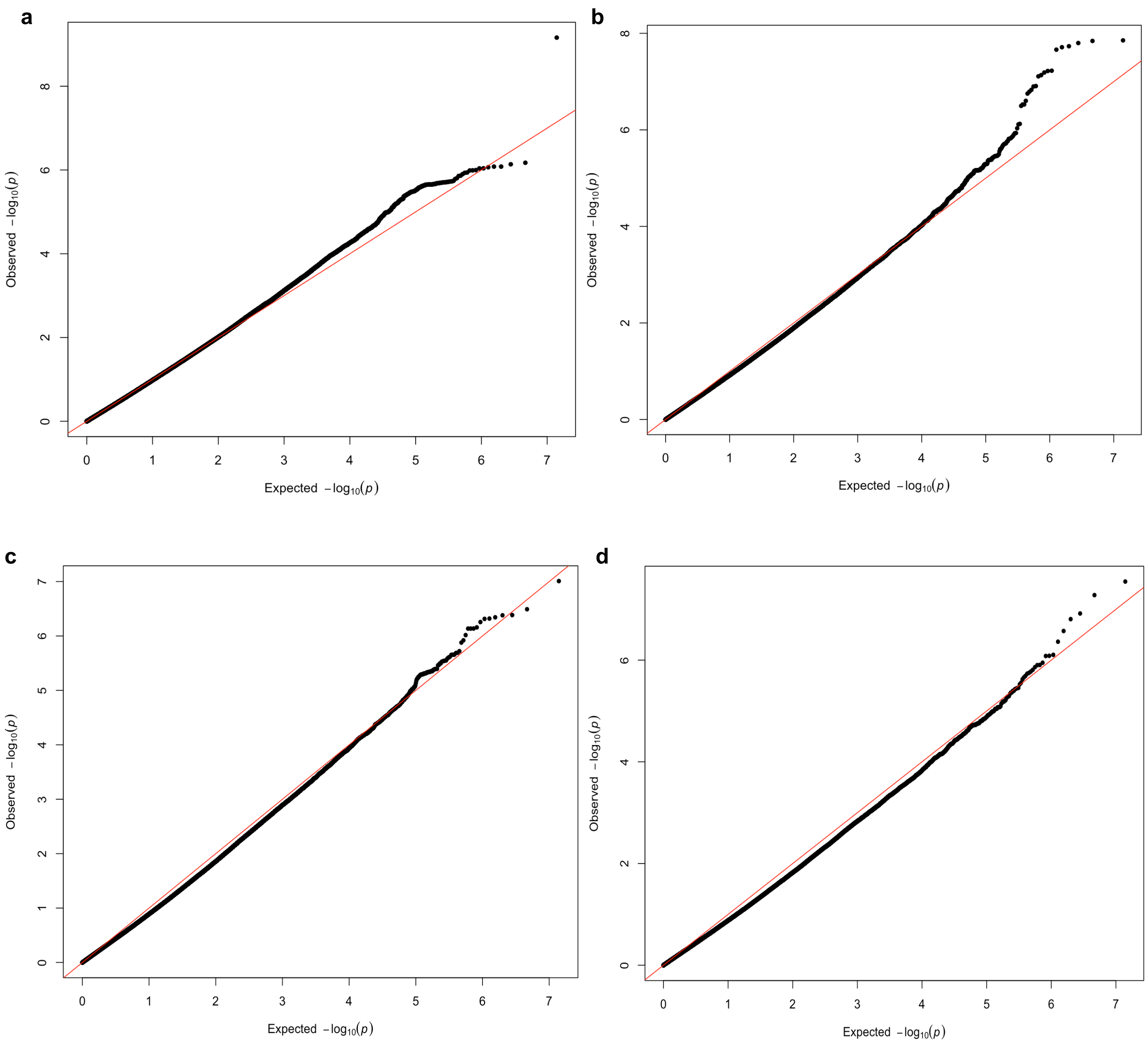


**Supplemental Figure 18. Genotype-by-time QQ plots.** Infant length (a), BMI (b), math (c), and reading (d). In (a) and (b), measures were taken at 6 , 8, and 12 months. There is a slight deflation in p-values. The traits were residualized on the full GRM prior to association testing. Leave-one-chromosome-out (LOCO) was not performed for computational efficiency and to account for the dense interfamilial relations in the sample. Note that genetic principal components (PCs) were included in the association testing step to control for population structure and other environmental confounding. These plots accompany Supplemental Figure 17.

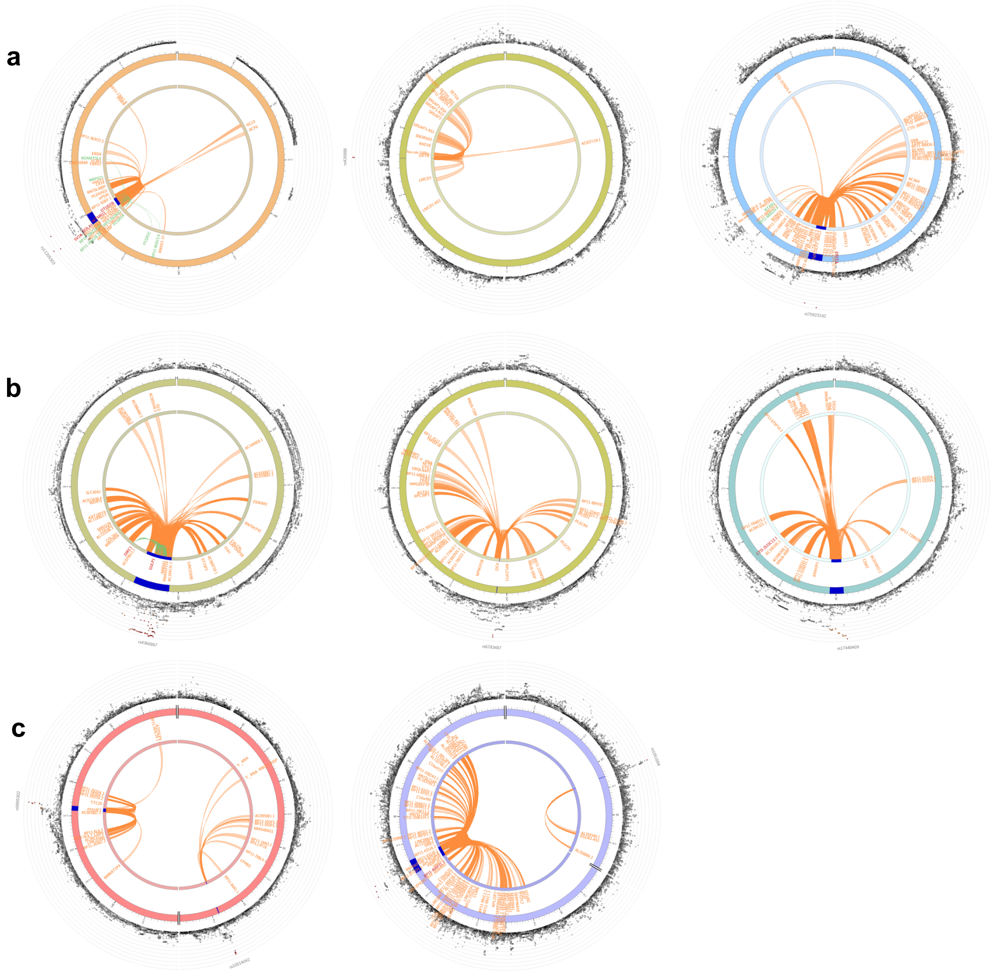

**Supplemental Figure 19. Intra-chromatin interactions of top non-additive loci.** Infant length includes top additive locus with time-varying effect: rs1205303 (a), math (b), reading (c). Plots are organized following Figure 2 in the main text. In (b, c), only non-additive loci that survived inverse normal transformation are shown. Dots in the outermost ring are SNPs plotted based on physical position along the chromosome, with red dots representing significant loci. The blue bar along the second ring represents the region found in Hi-C assays that interact with promoter or enhancer region of distal genes. The innermost ring is used to situate the genes along the chromosome. Orange = intra-chromatin interaction, green = eQTL, red = both. These plots are directly downloaded from FUMA.

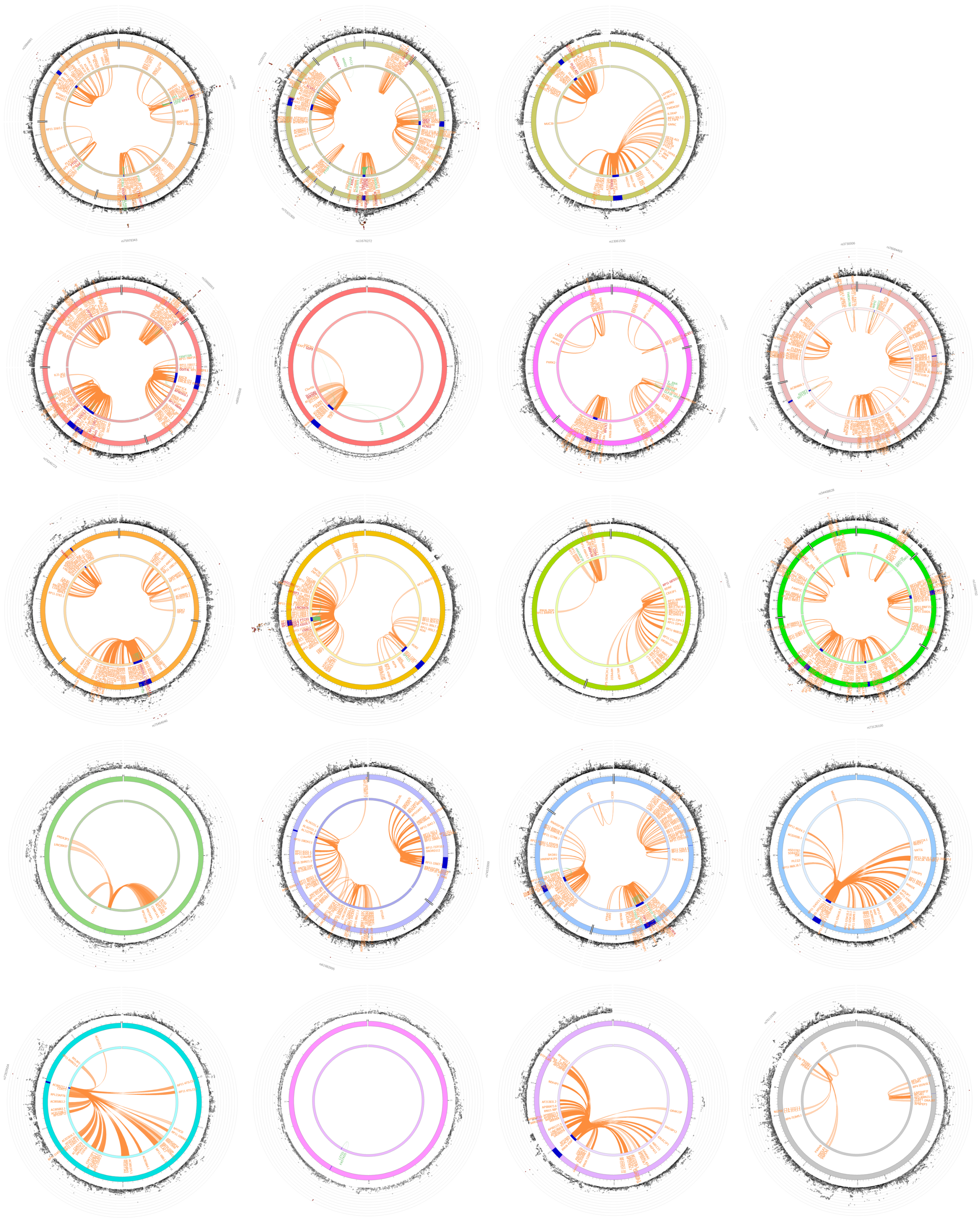

**Supplemental Figure 20. Intra-chromatin interactions of top BMI loci.** Plots are organized following Figure 2 in the main text. Dots in the outermost ring are SNPs plotted along the chromosome, with red dots representing significant loci. The blue bar along the second ring represents the region found in Hi-C assays that interact with promoter or enhancer region of distal genes. The ticks represent the position in million base pairs. The innermost ring is used to situate the genes along the chromosome. Orange = intra-chromatin interaction, green = eQTL, red = both. These plots are directly downloaded from FUMA.

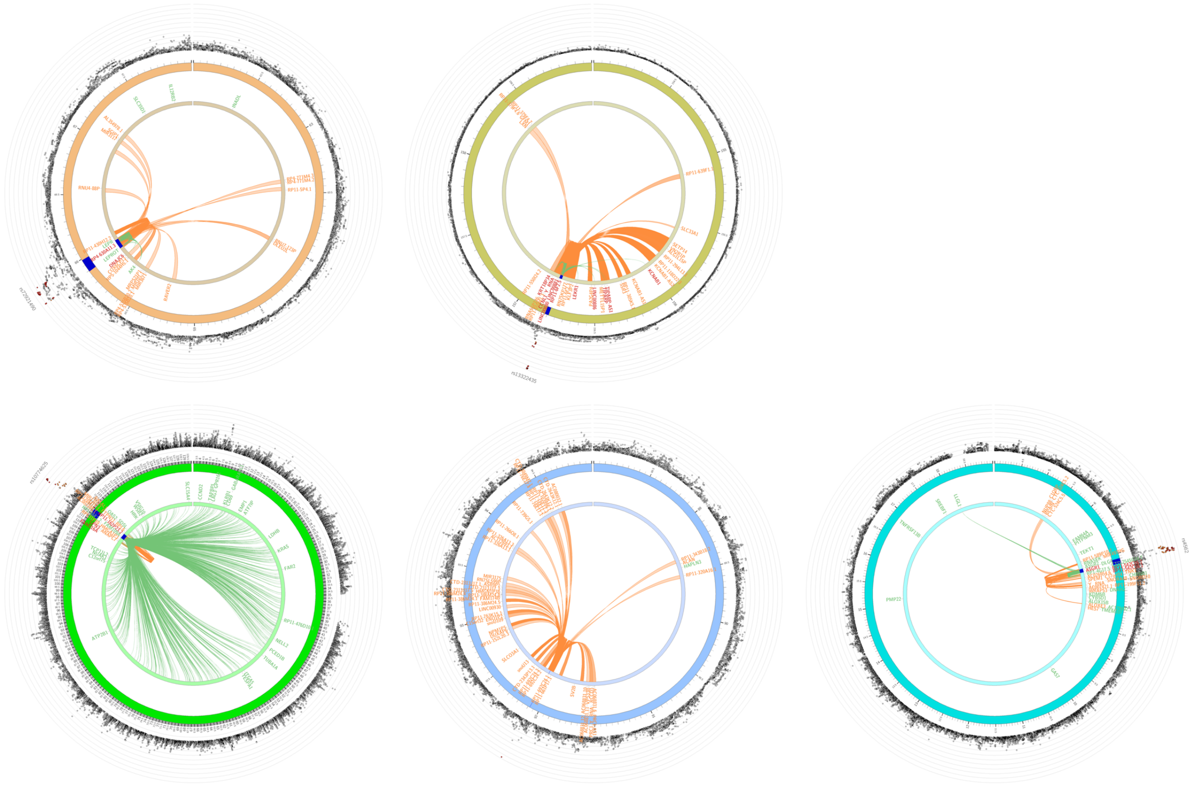

**Supplemental Figure 21. Intra-chromatin interactions of BMI loci with time-varying effects.** Plots are organized following Figure 5b in the main text. Dots in the outermost ring are SNPs plotted along the chromosome, with red dots representing significant loci. The blue bar along the second ring represents the region found in Hi-C assays that interact with promoter or enhancer region of distal genes. The ticks represent the position in million base pairs. The innermost ring is used to situate the genes along the chromosome. Orange = intra-chromatin interaction, green = eQTL, red = both. These plots are directly downloaded from FUMA.
